# Sequence and Structural Determinants of Ligand-dependent Alternating Access of a MATE Transporter

**DOI:** 10.1101/773572

**Authors:** Kevin L. Jagessar, Derek P. Claxton, Richard A. Stein, Hassane S. Mchaourab

## Abstract

MATE transporters are ubiquitous ion-coupled antiporters that extrude structurally- and chemically-dissimilar molecules and have been implicated in conferring multidrug resistance. Here, we integrate Double Electron Electron Resonance (DEER) in conjunction with functional assays and site-directed mutagenesis of conserved residues to illuminate principles of ligand-dependent alternating access of PfMATE, a proton-coupled MATE from the hyperthermophilic archaeon *Pyrococcus furiosus*. Pairs of spin labels monitoring the two sides of the transporter reconstituted into nanodiscs reveal large amplitude movement of helices that alter the orientation of a putative substrate binding cavity. We found that acidic pH favors formation of an inward-facing (IF) conformation, whereas elevated pH (>7) and the substrate rhodamine 6G stabilizes an outward-facing (OF) conformation. PfMATE isomerization between outward-facing and inward-facing conformations is driven by protonation of a previously unidentified intracellular glutamate residue that is critical for drug resistance. Our results can be framed in a mechanistic model of transport that addresses central aspects of ligand coupling and alternating access.

---

Bacterial homeostasis and survival is critically dependent on defense responses that modify, deactivate, or extrude cytotoxic molecules such as antiseptics and antibiotics, which passively cross the membrane down their concentration gradients^1^. One ubiquitous and highly conserved response entails the expression of polyspecific membrane transporters, referred to as multidrug (MDR) transporters, which harness the Gibbs energy stored in ion electrochemical gradients to power the uphill vectorial clearance of a broad spectrum of structurally- and chemically-dissimilar cytotoxic molecules^2–4^. The current dogma of active transport posits the energy-coupled isomerization of the transporter between multiple intermediates resulting in alternating access of the substrate binding site from one side of the membrane to the other^5, 6^. Defining the structural elements mediating alternating access and decoding the mechanism of energy conversion in a lipid bilayer environment are critical for a mechanistic description of coupled substrate efflux by MDR transporters.

Among the four families of ion-coupled MDR transporters in prokaryotes, the Multidrug and Toxin Extrusion (MATE) family includes Na^+^ and H^+^-coupled efflux transporters that are found in all three kingdoms of life^7–13^. Classified into three phylogenetic branches, the NorM, DinF, and Eukaryotic subfamilies, MATE transporters also share structural similarity with the multidrug/oligosaccharidyl-lipid/polysaccharide (MOP) transporter superfamily^14, 15^. Crystal structures of representatives of the three MATE branches revealed a conserved overall fold consisting of twelve transmembrane helices (TM) arranged in a unique topology as two bundles of six TMs and referred to hereafter as the N-lobe (TM1–TM6) and the C-lobe (TM7–TM12)^14^.

Until very recently, the canon of MATE structures was limited to outward-facing (OF) conformations in which the two lobes, related by an intramolecular two-fold symmetry and connected by a cytoplasmic loop, diverge and orient away from each other near the middle of the membrane forming a large central cavity open to the extracellular side^16–23^. Crystal structures of a NorM transporter from *N. gonorrheae*, NorM-Ng, bound to tetraphenylphosphonium, ethidium, and rhodamine 6G (R6G) demonstrated that substrates can bind to the same location within the central cavity between the two lobes^18^. Independently, we found that a spin labeled daunorubicin analog binds at a similar location in NorM from *V. cholera* (NorM-Vc)^24^. In contrast, a crystal structure of the proton-coupled DinF transporter PfMATE bound to a norfloxacin derivative suggested the docking of substrates to a distinct cavity within the N-lobe^19^.

Alternating access models postulate the isomerization of the transporter to an inward-facing (IF) state to which substrate binds and/or ions are released to the intracellular side. In the OF MATE structures, the putative central substrate binding cavities are shielded from the cytoplasm by highly ordered and packed protein regions, suggesting that transition to an IF conformation requires extensive structural rearrangements. A view of these rearrangements was recently captured from an IF crystal structure of PfMATE^25^. This structure, determined in the presence of native *P. furiosus* lipids, showed a change in the orientation of the central cavity that exposes its lumen to the intracellular side. Compared to the OF structure, TMs (2-6) and (8-12) in the two lobes undergo relative rigid body movement that disrupts helical packing and lead to the formation of new contacts. As highlighted in Fig. 1, a rupture of the interface between TMs 2 and 8 coupled with unwinding of TM1 repack the intracellular side of the transporter resulting in an IF cavity.

**Figure 1:**
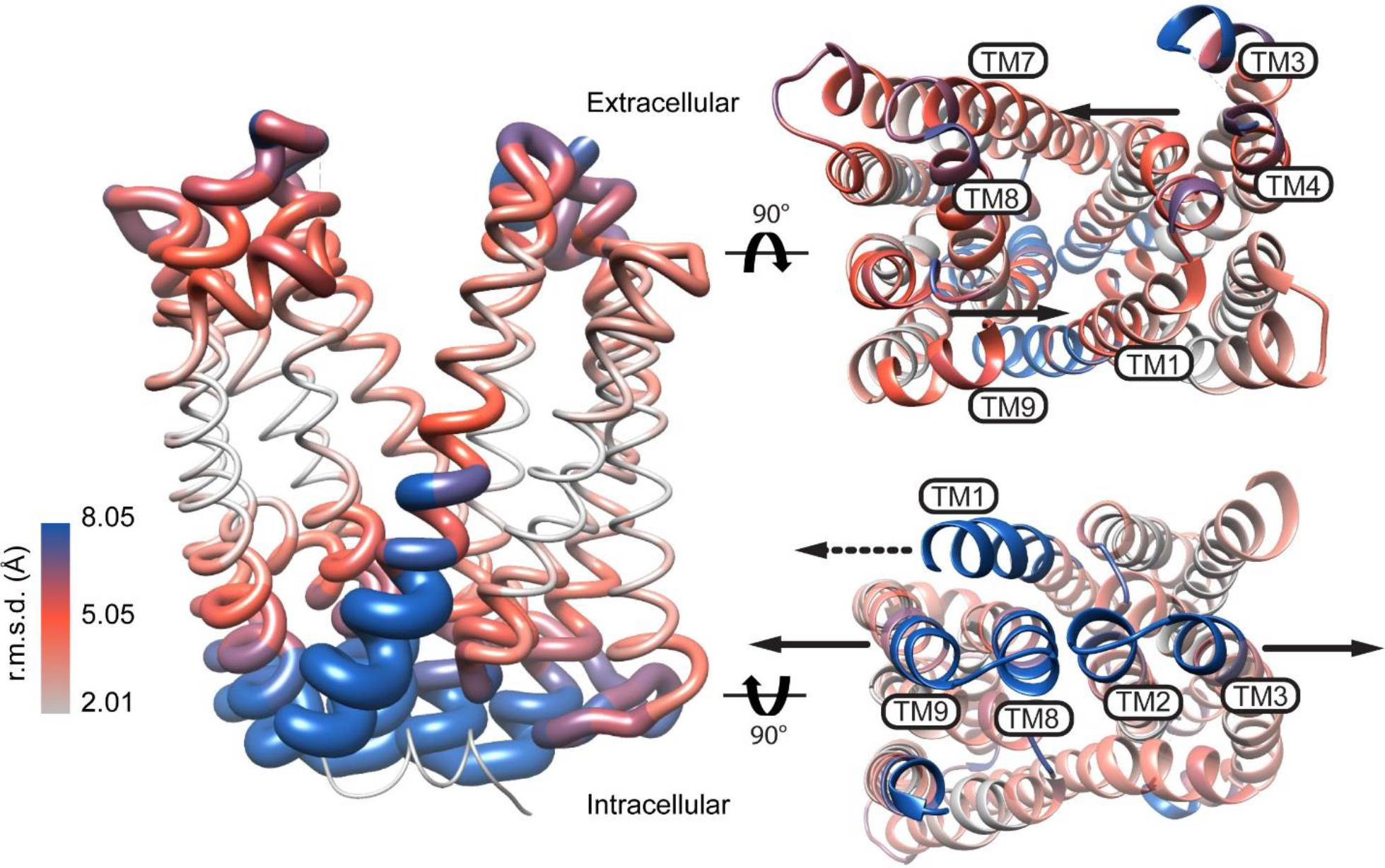
Model of PfMATE alternating access inferred from the crystal structures. The RMSD was derived from the alignment of the OF (PDB ID: 3VVN) and IF (PDB ID: 6FHZ) structures and depicted on a ribbon representation of the OF structure. Viewed from the extracellular and intracellular sides of the transporter, arrows show the direction of predicted movement of TMs in the OF to IF transition. The dashed arrow in the intracellular view points to TM1 unwinding.

Although the OF and IF structures of PfMATE suggest a blueprint of alternating access (Fig. 1), critical elements of the transport mechanism remain unresolved. Substrate/ion antiport entails differential stability of the conformations when either ligand is bound, yet Zakrzewska et al.^25^ found that the OF also crystallizes at low pH (pH 5-6.5), albeit in the absence of native lipids. Further confounding the mechanistic interpretation is the observation that structures of substrate- and ion-bound NorM-Ng as well as PfMATE at pH 8 and 6.0 were outward-facing, with bending of TM1 reported in the latter at lower pH^18, 19^. Finally, the residues and structural elements that couple ion gradients to conformational changes are not defined, although a network of conserved charged residues in the N-lobe of PfMATE was indirectly implicated^26^. Moreover, despite the lack of direct evidence for Na^+^ coupling, there has been speculation that Na^+^ rather than H^+^ may drive PfMATE alternating access^27^.

Prior to publication of the IF structure, we initiated a systematic Double Electron-Electron Resonance (DEER) ^28–32^ investigation of PfMATE to map proton- and substrate-dependent conformational changes in a lipid bilayer-like environment and to identify sequence motifs of ion and substrate coupling. For this purpose, an extensive network of spin label pairs was introduced at the extracellular and intracellular sides to interrogate ligand-dependent movements of TM helices. Here, we report that patterns of experimental distance distributions from DEER analysis reveal that an IF conformation is populated at pH 4 whereas pH 7.5 and substrate binding favor an OF conformation, demonstrating that protonation drives alternating access. Although these conformational changes were exclusively observed in lipid bilayers, native *P. furiosus* lipids were not required. Systematic mutagenesis of conserved residues uncovered an essential role for a previously unidentified residue, E163, in driving the proton-dependent isomerization of PfMATE. Together these findings can be integrated into a model of alternating access with mechanistic implications distinct from the crystal structures.

## Results

In the classic model of antiport, coupled ion/substrate transport entails the population of at least two conformational states, OF and IF, that are differentially stabilized by ligands. To avoid shorting the ion gradient, isomerization between the two states occurs only if one of the ligands is bound to the transporter. Therefore, to uncover the ligand dependence of PfMATE, which couples proton translocation to the cytoplasm to substrate extrusion to the periplasm, DEER distances distributions were determined at pH 4 to mimic a protonated state, at pH 7.5 to favor deprotonation, and in the presence of R6G at pH 7.5 to populate a substrate-bound state.

### Structural and functional integrity of PfMATE mutants

The functional profiles of the PfMATE mutants were analyzed by a recently described three-assay protocol^26^. First, we tested if the expression of the double cysteine mutants in *E. coli* conferred resistance to otherwise toxic concentrations of the drug R6G. Similar to WT-PfMATE, cells harboring the double cysteine mutants survived exposure to R6G concentrations that are lethal to cells transformed with empty vector (Supplementary Fig. 1), consistent with the mutants being functional in drug extrusion. Two pairs displayed resistance that was 20% of WT suggesting compromised activity as a consequence of the mutations. However, these mutants were not critical for our spectroscopic interpretation.

Second, we used fluorescence anisotropy to quantitatively measure the affinity of PfMATE spin labeled mutants to R6G *in vitro.* The apparent K_D_ indicated that spin labeled mutants bind the substrate with similar affinity as the WT (Supplementary Table 1).

Finally, the integrity of the proton conformational “switch” was assessed by monitoring pH-dependent Trp quenching. We have shown previously that protonation of WT-PfMATE leads to a reduction in Trp fluorescence, primarily from W44, reflecting localized structural rearrangements of TM1^26^. Robust Trp quenching for the spin labeled mutants at pH 4 (Supplementary Fig. 2) demonstrated H^+^-induced movement of TM1. As expected, mutants involving W44 displayed attenuated quenching in response to low pH. Taken together, these assays indicated that the cysteine mutations and subsequent spin labeling did not result in detectable structural or functional perturbations.

### Lipids are required for PfMATE conformational changes

Although DEER investigations of the MATE homolog NorM-Vc showed Na^+^ and H^+^-dependent conformational changes in β-dodecyl maltoside (DDM) micelles^33^, lowering the pH or addition of Na^+^ (data not shown) failed to induce substantial distance changes in DDM-solubilized PfMATE. Spin label pairs monitoring the intracellular and extracellular sides had similar distributions at pH 4 and 7.5 (Fig. 2, left panels), indicating the absence of large-scale conformational changes. In contrast, PfMATE reconstituted into nanodiscs composed of *E. coli* polar lipids and egg phosphotidylcholine (see methods) displayed evidence of large-scale distance changes upon protonation (Fig. 2, right panels). The strict lipid dependence of the pH-induced conformational changes is in agreement with the reported requirement of *P. furiosus* lipids for crystallization of the IF conformation. However, the DEER distance changes were observed in non-native lipid components. Furthermore, neither R6G binding at pH 7.5 nor addition of 50 mM Na^+^ at pH 7.5 elicited distance changes (Supplementary Fig. 3), pointing to protonation as the primary trigger of PfMATE isomerization in lipid bilayers.

**Figure 2:**
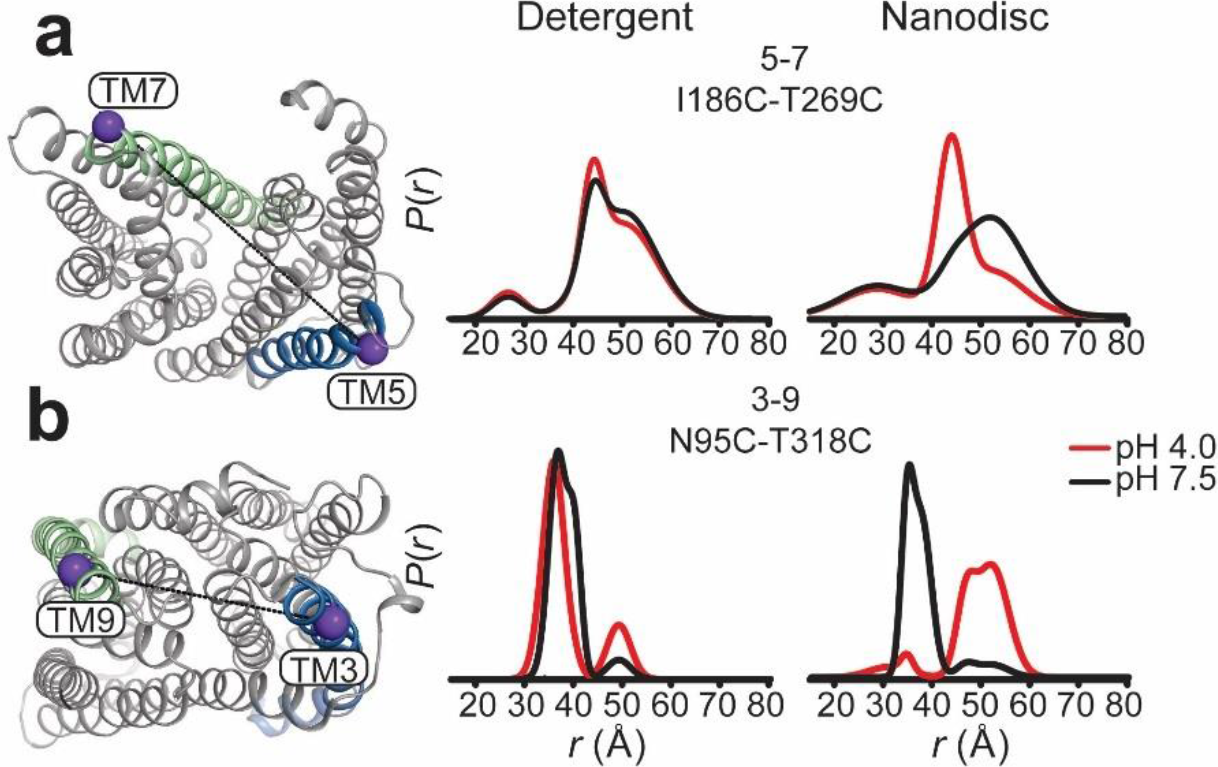
Ligand-dependent conformational dynamics of PfMATE requires a lipid environment. (**a**) Representative spin label pairs sampling distances between TM5 and TM7 on the extracellular side and (**b**) TM3 and TM9 on the intracellular side of PfMATE. The spin label locations are highlighted on the OF structure by purple spheres connected by a line. The helices targeted in the N-lobe and C-lobe are highlighted in blue and green, respectively. Distance distributions, representing the probability of a distance P(r) versus the distance (r) between spin labels, are shown in black traces at pH 7.5 and red traces at pH 4.0 in DDM micelles (left panel) and lipid nanodiscs (right panel).

### Protonation induces closing of the extracellular side

To determine the extent and amplitude of the pH-dependent conformational changes, two sets of spin labeled pairs were designed to survey the extracellular side of PfMATE. One set of pairs monitored distances between helices from the N-lobe to helices in the C-lobe (Figs 3-4). The other set consisted of labels monitoring distances between helices within each lobe (Supplementary Fig. 4).

In contrast to limited intradomain distance changes (Supplementary Fig. 4), a strikingly simple overall pattern emerged from the shifts in the distributions between the N- and C-lobe to shorter distances upon lowering the pH from 7.5 to 4 (solid black and red traces, respectively, in Figure 3). This pattern is highlighted by the relative movements that close the central cavity between TMs 7, 8 and 9 in the C-lobe and TMs 3, 4, 5 and 6 in the N-lobe. Although spin label pairs involving TM4 and TM11 were not as widely distributed, the observed distance changes are congruent with a relative movement between the N- and C-lobes. Unlike most TMs in the N-lobe, we did not observe distance changes between TM1 and TMs 7 and 8 suggesting movement of TM1 that is coupled to rearrangements of these helices in the C-lobe (Fig. 4).

**Figure 3:**
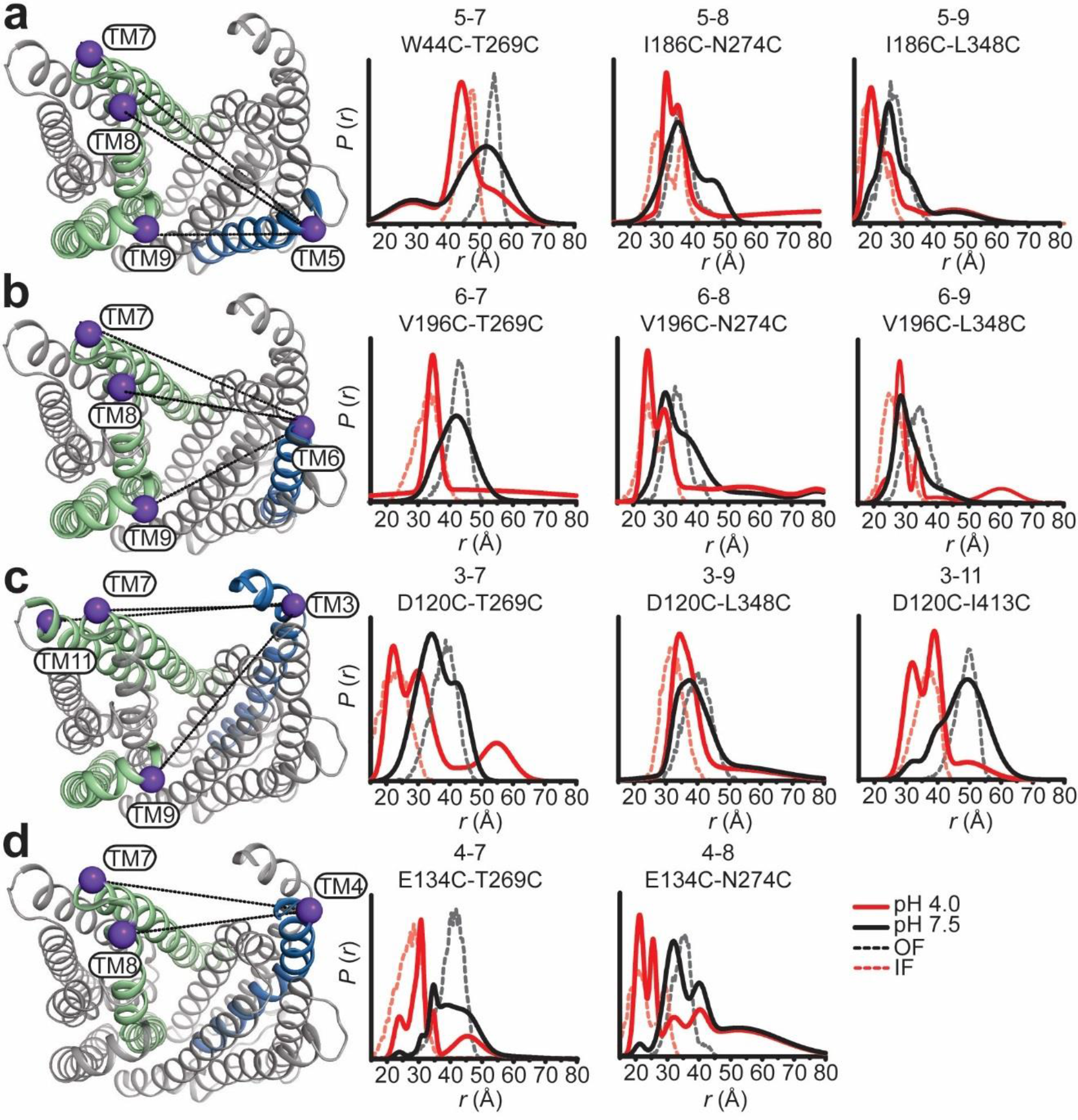
Protonation closes the extracellular side of PfMATE. Spin label pairs across the N- and C-lobes for DEER distance measurements are depicted on the extracellular side of the OF structure (**a** – **d**). Experimentally-determined distributions (solid lines) are plotted with the predicted distance distributions derived from the OF (black, dashed traces) and the IF (red, dashed traces) crystal structures. Measurements from TMs 7, 8, and 9 in the C-lobe to TMs 5 and 6 (**a** and **b**) in the N-lobe report decreased distances at pH 4 consistent with movement of these helices toward each other. Commensurate distance changes at pH 4 measured from TMs 3 and 4 in the N-lobe to the C-lobe (**c** and **d**) indicate closure of the extracellular side.

**Figure 4:**
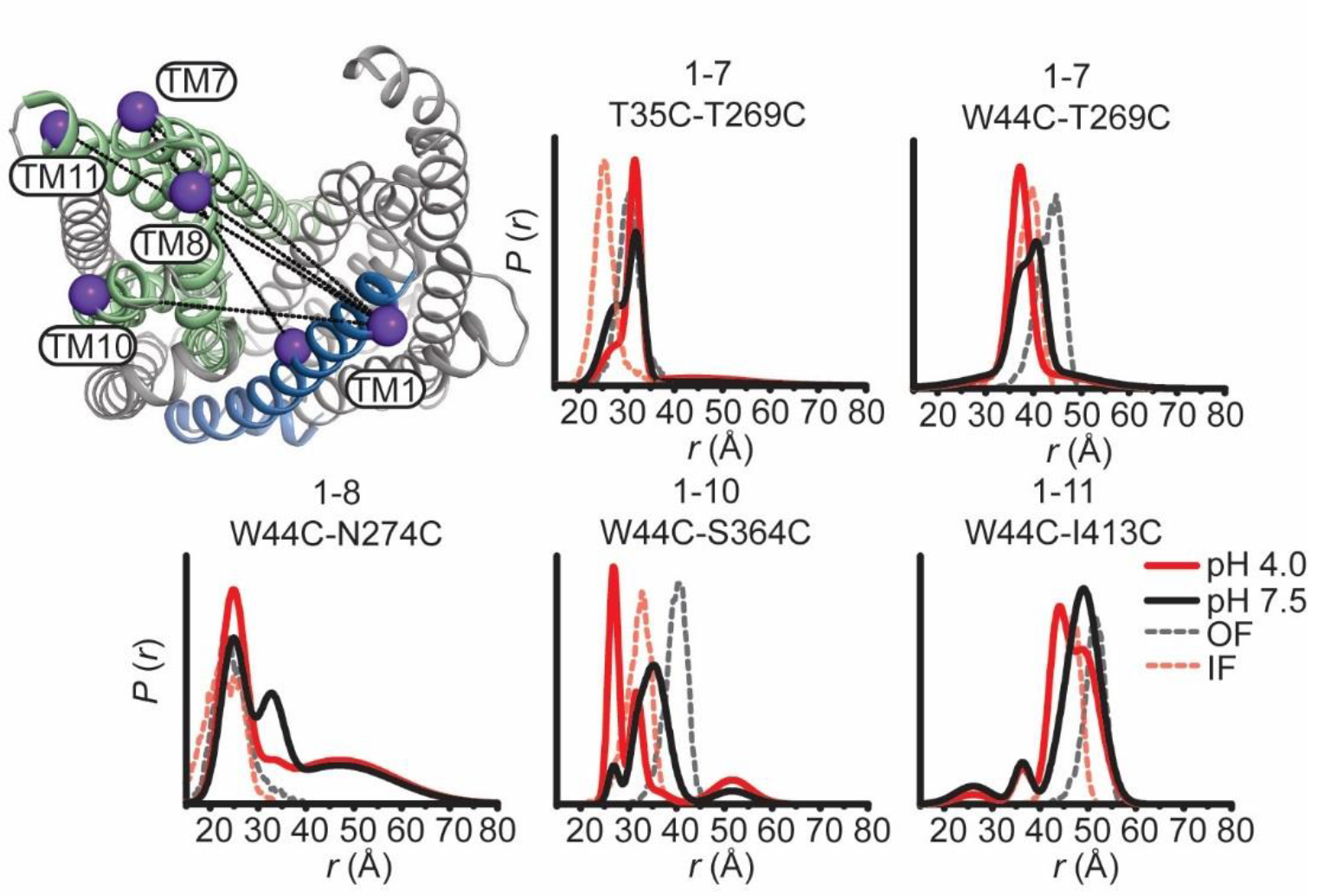
Movement of TM1 on the extracellular side of PfMATE is limited. The labeled positions on TM1 (purple spheres) to positions in the C-lobe for DEER measurements at pH 7.5 and pH 4 are depicted on the extracellular side of the OF structure. Distance distributions at pH 4 for TM1 are incongruent with predicted distributions based on the IF structure.

To determine if these structural changes are consistent in magnitude and direction with those expected based on the IF and OF conformations, we compared distance distributions predicted from two crystal structures (see methods, dashed distributions in Figures 3 and 4) with the experimental distributions. Except for TM1, we found that distributions calculated from the IF structure (6FHZ) overlapped with the pH 4 distributions whereas those calculated from the OF structure (3VVN) corresponded to the pH 7.5 distributions. This remarkable agreement between predictions and experiments link the structural rearrangements observed in the IF structure to protonation. It is also consistent with the canonical model of antiport wherein driving ions are expected to stabilize the IF conformation.

Notably, TM1 distance distributions to TMs 7 and 8 showed disagreements with the predicted distributions in both magnitude and direction (Fig. 4). A complex pattern of distance changes monitoring relative movement between TM1 and TM7 from two spin label pairs (T35C/T269C and W44C/T269C) reported distinct H^+^-dependent rearrangements, which may reflect twisting of TM1. Moreover, distance changes for TM1 were opposite to those predicted by the pH 6.0 OF crystal structure depicting a bent conformation of TM1^34^ (Supplementary Fig. 5). Although controversial^25, 35^, this conformation of TM1 was also captured in a crystal structure of a H^+^-coupled MATE from *V. cholerae* suggesting a conserved element of alternating access^23^. Therefore, the bent conformation of TM1 may reflect a structural intermediate along the pathway to the IF state.

### Protonation induces opening of the intracellular side

Coupled to the closing of the large central cavity on the extracellular side, protonation induced large amplitude movement of TM helices on the intracellular side (Fig. 5). Distributions between the N- and C- lobes shift to larger average distances at pH 4 relative to pH 7.5, in stark contrast to the extracellular side where distances predominantly decrease. The pattern of distance changes identified TMs 3 and 9 as focal points of conformational changes. TM3 moves away from TMs 1, 9 and 11 while TM9 moves away from TMs 6, 7 and 11 (Fig 5A, B, respectively).

**Figure 5:**
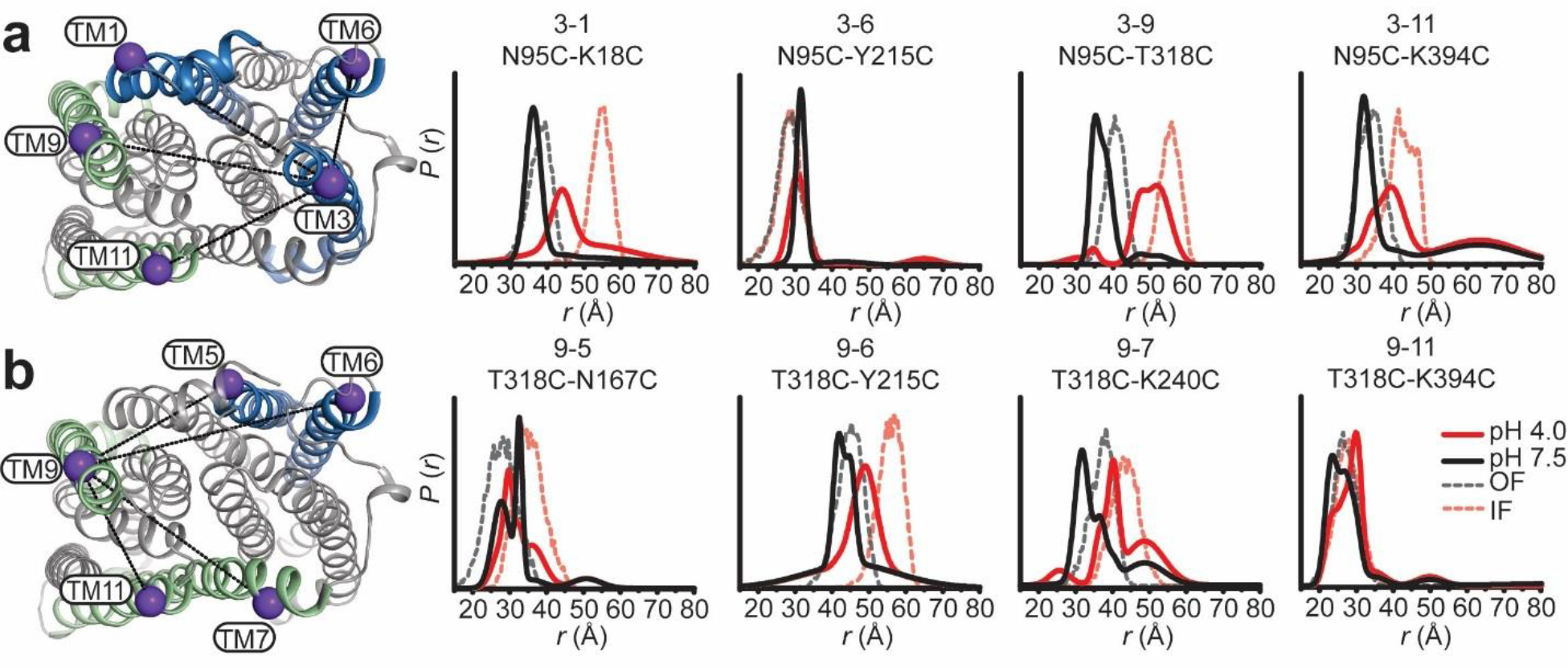
Relative movement of TM3 and TM9 on the intracellular side induced by protonation. Labeled positions for DEER measurements at pH 7.5 and pH 4 on TM3 and TM9 (**a** and **b**, respectively) to other TMs in the N- and C-lobes are depicted on the intracellular side of the OF structure. Protonation favors an increase in distance between the N- and C-lobes as predicted by the IF structure. The data identify TM3 and TM9 as foci of conformational changes, with an ∼ 15 Å increase in distance between these two helices at pH 4.

Except for TM1, we found a remarkable correspondence between the helices identified from pH-induced distance changes and those implicated in the opening of the intracellular side from comparison of the crystal structures (Figs. 5, 6). Moreover, predicted and experimental distance distributions partially overlap and the directions of the distance changes are identical, demonstrating that the intracellular side of the low pH conformation observed by DEER has similar features to the IF crystal structure.

Prominent discrepancies between predicted and experimental distributions are noted for the intracellular side of TM1, which in the IF structure unwinds leading to distance changes relative to TMs 3, 6, 11 and 12 (Figs. 5A, 6A). The experimental distance change between TM1 and TM3 was smaller than predicted, and distance distributions were almost superimposable at low and high pH between TM1 and TMs 6, 11 and 12. Together, these observations suggest that the intracellular side of TM1 remained intact under our conditions.

**Figure 6:**
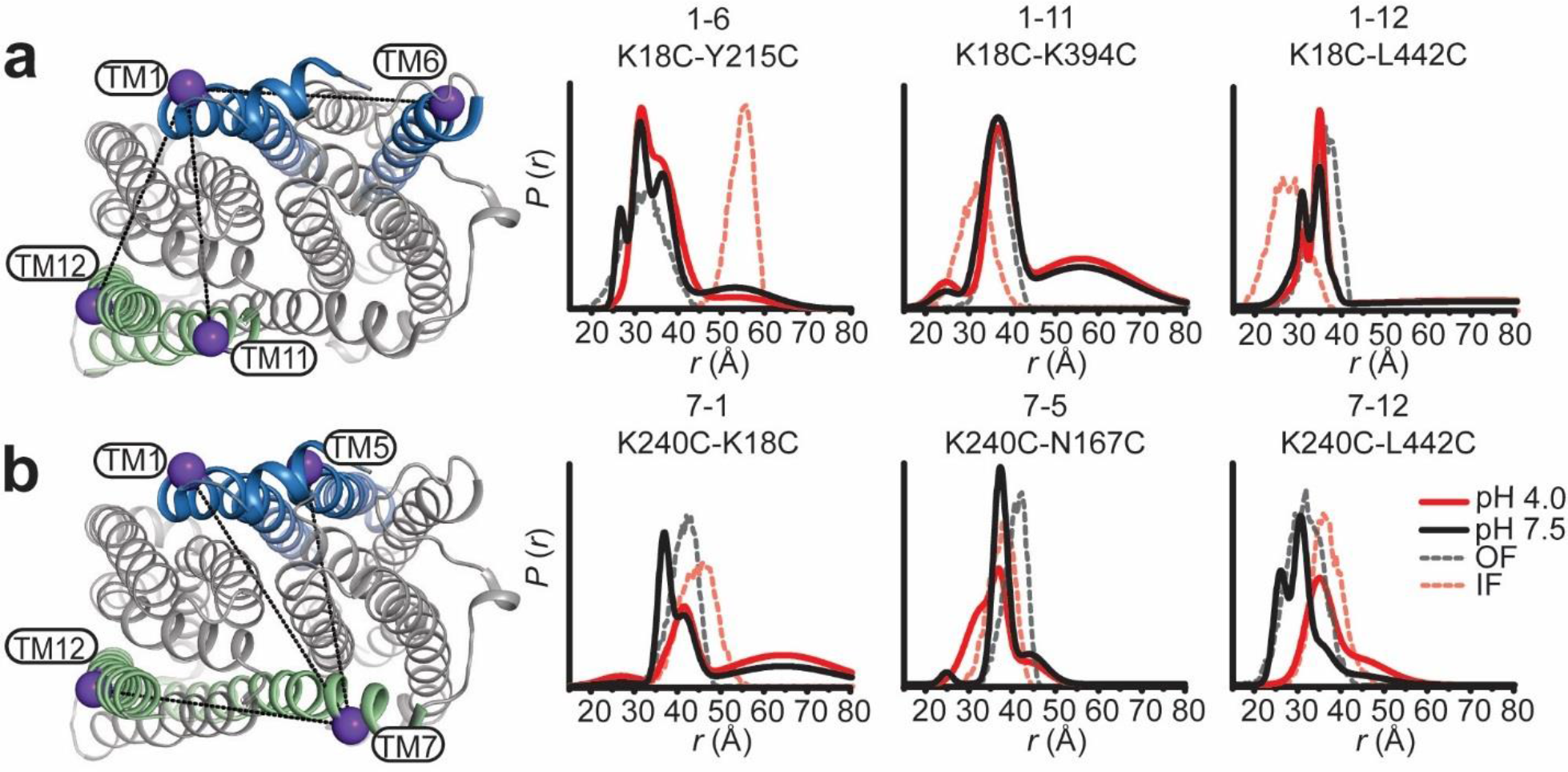
H^+^-dependent conformational changes of TM1 and TM7 are limited on the intracellular side. Positions for DEER measurements from TM1 and TM7 to helices in the N- and C-lobes are indicated on the OF structure. Relative distance changes involving TM1 are incongruent with the distances predicted from the IF structure (**a**). Measurement of TM7 labeled pairs (**b**) report changes in distance at pH 4 that suggest movement away from the C-lobe (TM7-TM12) and toward the N-lobe (TM7-TM5).

### Identification of residues involved in the protonation switch

An antiport mechanism requires the energetically downhill translocation of either Na^+^ or H^+^ to drive the uphill transport of the substrate. Whereas the NorM and DinF subfamily putatively couple to different ions, there has been evidence supporting a critical role for a conserved acidic residue on the extracellular side of TM1, D41 in PfMATE, that engages a number of amino acids in a network of H-bonds in the N-lobe^26^ (Fig. 7A). Protonation of D41 was predicted to induce rearrangement of this network^34^.

**Figure 7:**
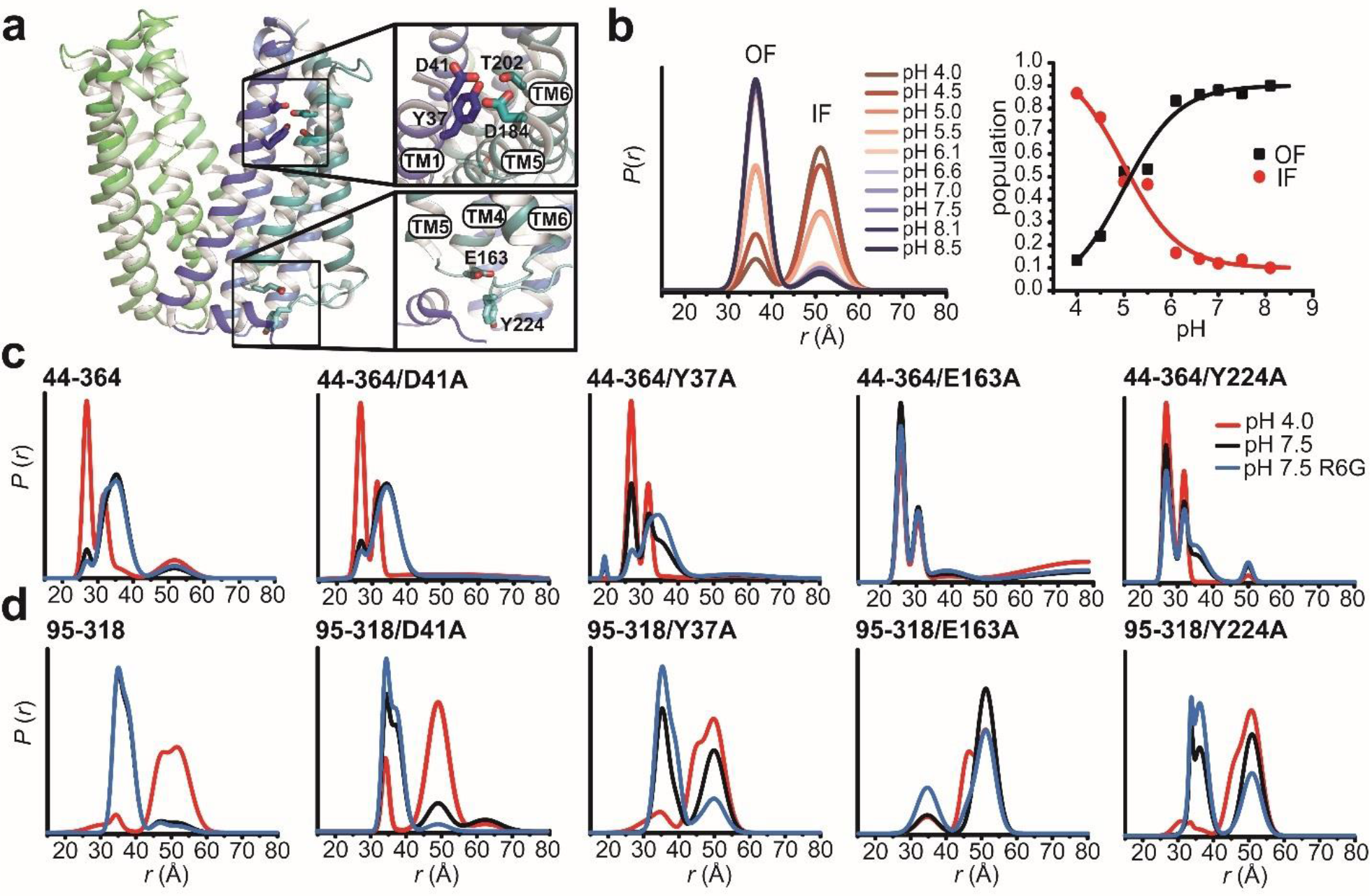
E163 is critical for the relative stability of the OF and IF conformations. Conserved residues previously identified in the N-lobe as crucial for PfMATE function are shown as sticks in the OF structure in the top inset (**a**). E163 on IL4-5 and Y224 on IL6-7 is depicted in the lower inset. Distance distributions acquired from titration of the intracellular reporter pair N95C-T318C from pH 4 to 8.5 were fit assuming a two-component Gaussian distance distribution (**b**, left panel) to quantify the variation in population as a function of pH (**b**, right panel). A pK of 5.1 was obtained from a non-linear least squares fit of the populations for the OF and IF peaks (**b**, black and red curves, respectively). Distance distributions of extracellular (W44C-S364C) and intracellular (N95C-T318C) reporter pairs (**c** and **d**, respectively, Supplementary Fig. 6) in acidic residue mutant backgrounds at pH 4, pH 7.5, and with R6G at pH 7.5 (blue traces). While D41A does not inhibit isomerization, Y37A, E163A, and Y224A abrogates the H^+^-dependent alternating access and destabilizes the OF conformation at pH 7.5.

To investigate if protonation of acidic residue(s) underpins the pH-dependence of alternating access, we determined the pK of the OF to IF transition. For this purpose, we measured the pH dependence of the distance change for the N95C/T318C intracellular pair monitoring relative movement of TM3 and TM9 (Fig. 7B). Global analysis of the distributions (see methods) yielded a titration curve depicting the populations of IF and OF (Fig. 7B) as a function of pH. A pK of approximately 5.1 was obtained from a non-linear least squares fit, which implicates the protonation/deprotonation of acidic residue(s) as the driver of PfMATE isomerization.

To pinpoint this residue or cluster of residues, we carried out mutagenesis of amino acids, including D41, that have been identified by previous studies as functionally critical and/or are highly conserved in MATE subfamilies^26, 34, 36^ (Fig. 7A, Supplementary Fig. 6A). OF/IF isomerization in these backgrounds was monitored by the TM1/TM10 extracellular pair W44C/S364C (Fig 7C) and the TM3/TM9 intracellular pair N95C/T318C (Fig. 7D). We found that substitutions of conserved residues D41A and D184A in the N-lobe cluster did not abrogate structural changes reported by the two pairs (Fig 7C, D; Supplementary Fig. 6B, C) although these substitutions compromised drug resistance and impaired pH-induced Trp quenching^26^ (Supplementary Figs 1, 2). Remarkably, Y37A was the only substitution in the N-lobe cluster that strongly attenuated coupled conformational changes (Fig. 7C, D). In the OF crystal structure, this residue interacts with D41 and D184 in an H-bond network (Fig. 7A) yet is unlikely to undergo protonation in the pH range identified above.

However, none of the mutations of residues in the N-lobe cluster fully mimicked the effects of alanine substitution of E163, located on an intracellular loop between TM4 and 5 (IL4-5) (Fig 7A). This substitution concomitantly inactivated PfMATE in R6G resistance (Supplemental Fig. 1) and abrogated conformational changes on both sides of the transporter (Fig. 7C, D). Moreover, distance distributions for this mutant shifted towards the protonated form at pH 7.5, suggesting that this glutamate is critical for the relative stability of the IF and OF conformations. To reinforce the importance of the negatively-charged sidechain, we mutated Y224, which is found on the cytoplasmic loop between TM6 and 7 (IL6-7) and in the OF structure is in close proximity to and potentially engages E163 in an H-bond or pi-charge interaction (Fig. 7A). Distance distributions for Y224A were shifted to favor the IF conformation at pH 7.5 on both sides of the transporter (Fig. 7C, D), supporting the notion that interaction with E163 modulates the relative stability of the IF and OF conformations. Thus, while a conserved network of polar and charged residues in the N-lobe is implicated in proton translocation, our data suggest that E163 is the protonation master switch that regulates the transition between IF and OF conformations.

### Substrate binding stabilizes the OF conformation

As noted above, distance distributions of spin label pairs in the WT background in the presence and absence of the drug R6G were superimposable at pH 7.5, suggesting that R6G binds to the OF conformation. To assess if the substrate drives the equilibrium towards the OF conformation, we took advantage of the Y37A, E163A, and Y224A mutations which reduce the relative stability of this conformation. We found that in the Y37A background, binding of R6G shifted the populations of the W44C/S364C and N95C/T318C reporter pairs toward the OF conformation at pH 7.5 (Fig. 7C, D), indicating that substrate binding stabilizes the OF conformation as would be expected for an antiporter. However, in the E163A and Y224A backgrounds, we found that binding of R6G shifted the populations of the intracellular N95C/T318C reporter pair but not of the extracellular W44C/S364C pair (Fig. 7D, C), possibly resulting in an occluded conformation and further underscoring the importance of E163 in OF/IF switching.

## Discussion

The extensive DEER analysis reported above illuminates principles of PfMATE alternating access, fills in critical gaps in the mechanism of substrate- and ion-coupling and sets the stage for understanding how lipids modulate the conformational cycle of the transporter. The remarkable agreement between experimental and predicted distance distributions, the latter calculated with modeling simplified dummy spin labels^37^, establishes that the DEER-detected, protonation-driven conformational changes in lipid nanodiscs describe, in outline, the OF to IF alternating access deduced from the crystal structures. Large amplitude distance changes, detected on both sides of PfMATE, reflect the pH-induced closing and opening of the extracellular and intracellular sides respectively. Although our results are consonant with a strict lipid dependence of PfMATE isomerization, we found that native lipids are not absolutely required. It is conceivable that endogenous lipids, predominantly consisting of diphytanyl acyl chains, may shape the energetics of alternating access in a native cellular environment.

### PfMATE isomerization is driven by protonation

Protonation, experimentally achieved by lowering the pH, is sufficient to support transition from OF to IF states in the presence of non-native lipids. Binding of R6G did not engender large conformational changes in a WT background. That the substrate stabilizes the OF conformation was exposed from distance measurements in mutant backgrounds that shift the equilibrium towards the IF conformation.

The original structures by Tanaka et al.^34^ purportedly identified a proton binding site in a cluster of conserved residues in the N-lobe. However, subsequent molecular dynamics simulations, in conjunction with reexamination of the crystallographic data, suggested that Na^+^ is bound at this site^27^. Although our findings do not weigh in specifically on this question, they conclusively demonstrate that Na^+^ binding does not drive the transition from OF to IF (Supplementary Fig. 3). Thus, the ligand dependence captured by the DEER analysis establishes that alternating access of PfMATE is proton-coupled.

### E163 is the master protonation switch

Reinforcing this conclusion is the finding that alanine substitution of a single acidic residue, E163, found on an intracellular loop between TM4 and 5 abrogates the proton-dependence of alternating access and destabilizes the OF conformation resulting in an IF conformation at pH 7.5. Specifically, disruption of the interaction between E163 and Y224 leads to a shift in distance populations indicating that the stability of the OF intermediate is compromised (Fig. 7C, D). Although Y224 is not strongly conserved, analysis of 500 homologs of PfMATE indicates that Glu is present at position 163 in 45% of the sequences (data not shown). Collectively, these results suggest that the specific interaction between IL4-5 and IL6-7 mediate the stability of the OF conformation, which may have implications across the MATE family (Supplementary Fig. 7). We propose that the large IL6-7 loop may function as a “belt” that regulates the conformational landscape reminiscent of the mechanism proposed for the MOP transporter MurJ^38^. In this mechanism, straightening of the intracellular portion of TM7 associated with formation of the OF in MurJ alters the position of the IL6-7 loop relative to the membrane and provides tension to support closure of the intracellular gate.

Previous investigations into residues involved in ion-coupling focused on the N-lobe cluster, such as D41, and to a lesser extent on residues in the C-lobe implicated in binding of Na^+^ congeners in the crystal structures of NorM transporters^9, 24, 33, 39–44^. Our previously published work and the data reported here support the involvement of the N-lobe residues in PfMATE function as reflected in compromised drug resistance and reduced Trp quenching, which has been interpreted as impaired formation of a H^+^-dependent structural intermediate required for the transport cycle^26^. However, alanine substitutions of these residues do not completely abrogate the OF/IF isomerization (Supplementary Fig. 6).

To resolve this apparent discrepancy, it is instructive to contrast the direct detection of conformational changes with resistance assays which do not report exclusively on transporter isomerization. Combined with the blunt nature of mutagenesis, reduced resistance could result from convoluted effects of local structural distortion, interference with ligand binding, and/or changes in transport kinetics. The latter is particularly confounding because the ability of a transporter to confer resistance depends on a delicate balance between the passive diffusion rate of hydrophobic substrates through the membrane bilayer and its rate of extrusion by the transporter. Thus, increasing the dimensionality of the analysis by combining direct detection of conformational dynamics with functional assays provides unique insight into the role of sequence and structural motifs in the transport mechanism.

### Model of PfMATE ligand dependent alternating access

Framed in the context of the available structures, our results lead to an antiport model of PfMATE that reveals how protons and substrate differentially modulate the stability of the IF and OF conformations, identifies residues critical for protonation, and suggests a putative path for proton translocation (Fig. 8). In this model, the stable resting state is OF (Fig. 8A) whereas the IF (Fig. 8C) is of relatively high energy. Binding of R6G stabilizes the OF whereas protonation drives the isomerization to the IF state (Fig. 8, B to C), as would be expected if the proton motive force powers alternating access.

**Figure 8:**
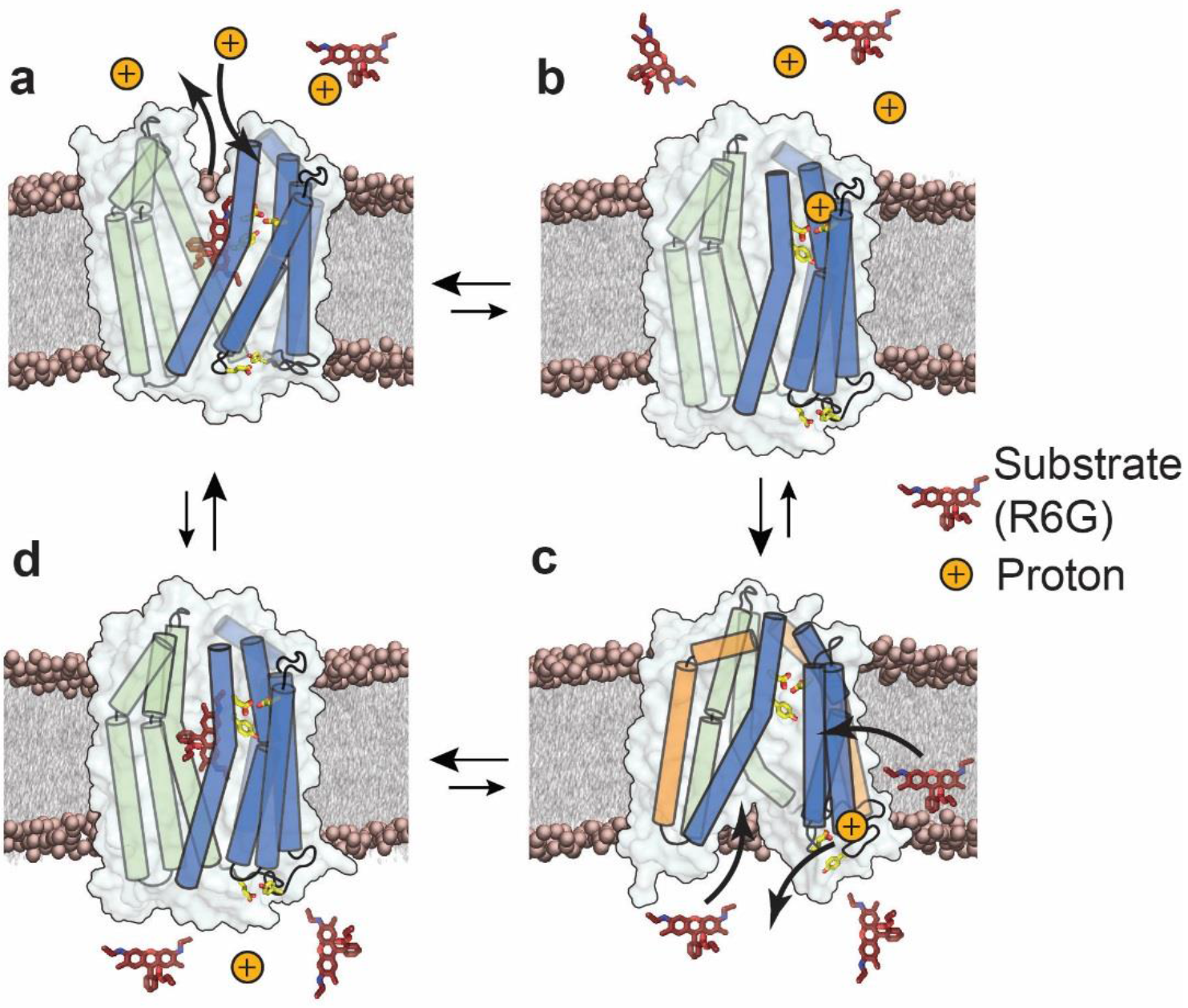
Proposed model of antiport for PfMATE. N- and C-lobe helices are depicted as cylinders and are colored blue and green, respectively. For clarity not all helices are shown. Side chains of Y37, D41 and D184 on the extracellular side, and E163 and Y224 on the intracellular side are represented as sticks. In the resting state, substrate is bound to the transporter and stabilizes the OF conformation (**a**), (Supplementary Fig. 3, black, blue traces; appendices). Upon protonation of residues in the N-lobe cluster, substrate is released and the transporter could isomerize to an occluded conformation (**b**). Proton translocates from the extracellular side to the intracellular side, where E163 undergoes protonation thereby disrupting the IL4-5/IL6-7 interaction and opening the intracellular side through movements of TM3 and TM9 (orange cylinders, **c**), (Figs 5, 6 red traces). The IF conformation binds drug, either from the cytoplasm or from the inner leaflet of the membrane bilayer (**c**), upon which the transporter isomerizes to a drug bound occluded conformation (**d**), (Fig. 7c, **d**) which in turn isomerizes spontaneously to the OF resting state (**a**).

We thus propose that the high energy IF state is destabilized by the terminal step of proton dissociation from E163. To ensure coupled antiport, proton dissociation must occur simultaneously with binding of substrate (Fig 8, C to D). The data presented here does not weigh in on whether the proton/substrate competition is direct or indirect. However, we have previously suggested that H^+^ and substrate occupy distinct binding sites^26^. Following proton release to the intracellular side, the transporter, bound to substrate, spontaneously isomerizes to the OF conformation (Fig 8, D to A).

Although primarily cast in terms of explicit OF/IF conformations, the model does not exclude the possibility of the population of a membrane accessible but solution-occluded conformations (Fig 8B, D) as we have proposed for LmrP and which have been presented for MurJ^38, 45^. Such a conformation may serve to enable substrate binding from the inner leaflet of the bilayer while excluding protons. This conformation should be of similar energy to the OF state in light of the marginal stabilization of the OF by the substrate R6G. The occupation of a substrate-bound doubly occluded state (Fig. 8C) may be inferred from the analysis of W44C/S364C and N95C/T318C where alanine substitution of critical residues (E163 and Y224) supported closing of the intracellular side but not concomitant opening of the extracellular side (Fig 7C-D).

A combination of approaches, including sequence analysis, spectroscopic studies, and molecular dynamics simulations, have underscored the significance of a conserved residue network in the N-lobe cavity to the transport mechanism^19, 26, 27, 33^. This cavity likely serves as the ion entry point^46^, but the ion translocation pathway to the intracellular side has not been defined experimentally. Further examination of the PfMATE sequence through alignments and conservation analysis (ConSurf server^47, 48^) outlines a putative pathway from the N-lobe cavity to the master switch, E163. This pathway is lined by a cascade of polar and charged sidechains, which may be capable of water-mediated proton transfer. The integrated approach described here will enable testing the roles of these residues in the alternating access mechanism of PfMATE.

## Methods

### Site-Directed Mutagenesis

Wild-type PfMATE was cloned into pET19b vector encoding an N-terminal 10-His tag under control of an inducible T7 promoter and was used as the template to introduce double cysteine pairs and background mutations via site-directed mutagenesis with complementary oligonucleotide primers. Substitution mutations were generated using a single-step PCR in which the entire template plasmid was replicated from a single mutagenic primer. PfMATE mutants were sequenced using both T7 forward and reverse primers to confirm mutagenesis and the absence of aberrant changes. Mutants are identified by the native residue and primary sequence position followed by the mutant residue.

### Expression, Purification, and Labeling of PfMATE

C43 (DE3) cells were freshly transformed with pET19b vector encoding wild type or mutant PfMATE. A transformant colony was used to inoculate Luria-Bertani (LB) media which was grown overnight (∼15 h) at 34 °C and was subsequently used to inoculate 3 L of minimal medium A at a 1:50 dilution. Cultures were incubated while being shaken at 37 °C until they reached an absorbance at 600 nm (Abs_600nm_) of ∼0.8, at which time the expression of PfMATE was induced by the addition of 1mM IPTG. The cultures were incubated overnight (∼15 h) at 20 °C and then harvested by centrifugation. Cell pellets were resuspended in 20 mL of resuspension buffer [20 mM Tris-HCl, pH 7.5, 20 mM NaCl, 30 mM imidazole, and 10% (v/v) glycerol], including 10 mM DTT, and lysed by five passes through an Avestin C3 homogenizer. Cell debris was removed by centrifugation at 9,000 × g for 10 min. Membranes were isolated from the supernatant by centrifugation at 200,000 × g for 1.5 h.

Membrane pellets were solubilized in resuspension buffer containing 20 mM β-DDM (Anatrace) and 0.5 mM DTT and incubated on ice with stirring for 1 hour. Insoluble material was cleared by centrifugation at 200,000 × g for 30 min. The cleared extract was bound to 1.0 mL Ni-NTA Superflow resin (bed volume) at 4 °C for 2 h. After washing with 10 bed volumes of buffer containing 30 mM imidazole, PfMATE was eluted with buffer containing 300 mM imidazole.

Double cysteine mutants were labeled with two rounds of 20-fold molar excess 1-oxyl-2,2,5,5-tetramethylpyrroline-3-methyl methanethiosulfonate (Enzo Life Sciences) per cysteine at room temperature in the dark over a 4-hour period, after which the sample was placed on ice at 4 °C overnight (∼15 h). Unreacted spin label was removed by size exclusion chromatography over a Superdex200 Increase 10/300 GL column (GE Healthcare) into 50 mM Tris/MES, pH 7.5, 1 mM β-DDM, and 10% (v/v) glycerol buffer. Peak fractions of purified PfMATE were combined and concentrated using a 100,000 MWCO filter concentrator (Millipore) and the final concentration was determined by A_280_ measurement (*ε* = 46870 M^−1^⋅cm^−1^) for use in subsequent studies.

### Reconstitution of PfMATE into Nanodiscs

*E. coli* polar lipids and PC (L-α phosphatidylcholine) (Avanti Polar Lipids, Alabaster, USA) were combined in a 3:2 w/w ratio, dissolved in chloroform, evaporated to dryness on a rotary evaporator and desiccated overnight under vacuum in the dark. The lipids were hydrated in 50 mM Tris/MES pH 7.5 buffer to a final concentration of 20 mM, homogenized by freezing and thawing for 10 cycles, and stored in small aliquots at −80 °C. MSP1D1E3 was expressed and purified as previously described and dialyzed into 50mM Tris/MES pH 7.5 buffer^49^. MSP1D1E3 was concentrated using a 10,000 MWCO filter concentrator (Millipore) and the final protein concentration was determined by A_280_ measurement (*ε* = 29,910 M^−1^⋅cm^−1^).

For reconstitution into nanodiscs, spin labeled double cysteine mutants in β-DDM micelles were mixed with *E. coli* polar lipids/PC lipid mixture, MSP1D1E3 and sodium cholate in the following molar ratios: lipid:MSP1D1E3, 50:1; MSP1D1E3:PfMATE, 10:1; cholate:lipid, 3:1. Reconstitution reactions were mixed at 4 °C for 1 hour. The detergent was removed from the reaction by addition of 0.1 g/ml Biobeads (Bio-Rad) and incubation at 4 °C for 1 hour. This was followed by another addition of 0.1 g/ml Biobeads with 1-hour incubation, after which 0.2 mg/ml Biobeads were added and mixed overnight. The next day, 0.2 mg/ml Biobeads were added and mixed for one hour^28^. The reaction was filtered using a 0.45 µm filter to remove Biobeads. Full nanodiscs were separated from empty nanodiscs by size exclusion chromatography into 50 mM Tris/MES, pH 7.5 and 10% (v/v) glycerol buffer. The PfMATE-containing nanodiscs were concentrated using Amicon ultra 100,000 MWCO filter concentrator, then characterized using SDS-PAGE to verify reconstitution and estimate reconstitution efficiency. The concentration of spin labeled mutants in nanodiscs was determined as described previously by comparing the intensity of the integrated CW-EPR spectrum to that of the same mutant in detergent micelles^50^.

### CW-EPR and DEER Spectroscopy

CW-EPR spectra of spin labeled PfMATE samples were collected at room temperature on a Bruker EMX spectrometer operating at X-band frequency (9.5 GHz) using 10 mW incident power and a modulation amplitude of 1.6 G. DEER spectroscopy was performed on an Elexsys E580 EPR spectrometer operating at Q-band frequency (33.9 GHz) with the dead-time free four-pulse sequence at 83 K^30^. The pulse lengths were 10-14 ns (π/2) and 20 ns (π) for the probe pulses and 40 ns for the pump pulse. The frequency separation was 63 MHz. To ascertain the role of H^+^, samples were titrated to pH 4 with an empirically determined amount of 1 M citric acid and confirmed by pH microelectrode measurement. The substrate-bound state was generated by addition of 1 mM R6G at high and low pH. Samples for DEER analysis were cryoprotected with 24% (v/v) glycerol and flash-frozen in liquid nitrogen.

Primary DEER decays were analyzed using home-written software operating in the Matlab (MathWorks) environment as previously described^49, 51^. Briefly, the software carries out global analysis of the DEER decays obtained under different conditions for the same spin labeled pair. The distance distribution is assumed to consist of a sum of Gaussians, the number and population of which are determined based on a statistical criterion. A noticeable change in the intermolecular background was observed in nanodiscs at pH 4 giving rise to a steep decay relative to pH 7.5. This change in background was reversible by returning to pH 7.5. Negative stain electron microscopy analysis of a related MATE transporter suggests that this pH-dependent change in background is associated with reversible clustering of individual nanodisc particles (data not shown). We also analyzed DEER decays individually and found that the resulting distributions are in agreement with those obtained from global analysis. The differences were primarily in the number of gaussian components required for the fit. Because our analysis assumes two conformations, IF and OF, the differences in the shape of the distance components is not material to our conclusion.

Distance distributions on the PfMATE crystal structures [Protein Data Bank (PDB) ID code 3VVN, 6FHZ] were predicted *in silico* using molecular-dynamics simulations with dummy spin labels (MDDS), which were facilitated by the DEER Spin-Pair Distributor at the CHARMM-GUI website^37^.

### R6G Resistance Assay

Resistance to R6G toxicity was carried out as previously described^26^. *Escherichia coli* BL21(DE3) were transformed with empty pET19b vector, pET19b encoding PfMATE wild-type, or mutant PfMATE. A dense overnight culture from a single transformant was used to inoculate 10 mL of LB broth (LabExpress) containing 0.1 mg/mL ampicillin (Gold Biotechnology) to a starting Abs_600nm_ of 0.0375. Cultures were grown to Abs_600nm_ of 0.3 at 37 °C and expression of the encoded construct was induced with 1.0 μM IPTG (Gold Biotechnology). Expression was allowed to continue at 37 °C for 2 h, after which the Abs_600nm_ of the cultures was adjusted to 0.5. The cells were then used to inoculate (1:20 dilution, starting Abs_600nm_ = 0.025) a sterile 96-well microplate (Greiner Bio-one) containing 50% LB broth, 0.1 mg/mL ampicillin, and 60-72 μg/mL R6G. Microplates were incubated at 37 °C with shaking at 200 rpm for 6-10 h, after which the cell density was measured on a BioTek Synergy H4 microplate reader and normalized to the wild type construct. Each data point was performed in triplicate, and the experiment was repeated to obtain the mean and standard error of the mean (S.E.M). To statistically determine the impact of substitutions on R6G resistance, a one-way ANOVA in the program Origin (OriginLab) determined that the difference between population means was statistically significant at the 0.05 level.

### Fluorescence Anisotropy

R6G was solubilized in water and dilutions in ethanol were prepared and measured spectrophotometrically at 524 nm (*ε* = 116,000 M^−1^⋅cm^−1^) to determine the concentration. Dilutions of purified PfMATE in 50 mM Tris/MES pH 7.5, 10% glycerol (v/v),1mM β-DDM buffer were mixed with a constant concentration of R6G (2.0 uM) in a total volume of 25 μL in a 384-well black fluorescence microplate (Greiner Bio-One) and incubated at room temperature for >5 min. R6G fluorescence anisotropy was measured using a BioTek Synergy H4 microplate reader with a 480 nm excitation filter (20 nm band pass) and a 570 nm emission filter (10 nm band pass)^52^. Each point was measured in triplicate and R6G binding affinity was determined by a non-linear least squares analysis of each individual curve in Origin. The average *K*_D_ and standard deviation (S.D.) for each mutant are reported.

### Tryptophan Fluorescence Quenching

Purified PfMATE in 50 mM Tris/MES pH 7.5, 10% glycerol (v/v)1mM β-DDM buffer was adjusted to pH 4.0 using an empirically determined volume of citric acid. Samples at pH 7.5 were adjusted with an equivalent volume of buffer to maintain an equal concentration of protein between pH conditions. Samples were placed in a quartz fluorometer cell (Starna Cells, Inc; cat #: 16.40F-Q-10/Z15) and tryptophan fluorescence quenching was monitored using a T-format fluorometer from Photon Technology International at 23 °C. The excitation wavelength was (λ_ex_) 295 nm and tryptophan fluorescence was monitored between (λ_em_) 310 nm and 370 nm. The fluorescence intensity at 329 nm was recorded from the spectra to determine the difference in fluorescence intensity between pH 7.5 and pH 4.0 samples^26^. Experiments were repeated in triplicate and the mean and S.E.M of fluorescence quenching was determined. A one-way ANOVA in Origin to assess the impact of substitutions on Trp quenching revealed that there was a statistically significant difference between population means at the 0.05 level.

## Acknowledgments

The authors wish to thank Dr. Reza Dastvan for modeling studies and helpful discussions regarding data analysis. We also wish to thank Dr. Smriti Mishra for help in optimization of conditions for nanodisc reconstitution and Ms. Abigail Poff and Mr. Nabeeh Daouk for their assistance in creating mutants and protein expression and purification. This work was supported by an NIH grant R01 GM077659 to H.S.M.

## Author Contributions

K.L.J, D.P.C, and H.S.M conceived and designed experiments. K.L.J and D.P.C performed experiments. K.L.J, D.P.C, and R.A.S. analyzed the data. H.S.M, K.L.J, and D.P.C wrote the paper.

## Supplementary Information

**Supplementary Figure 1:**
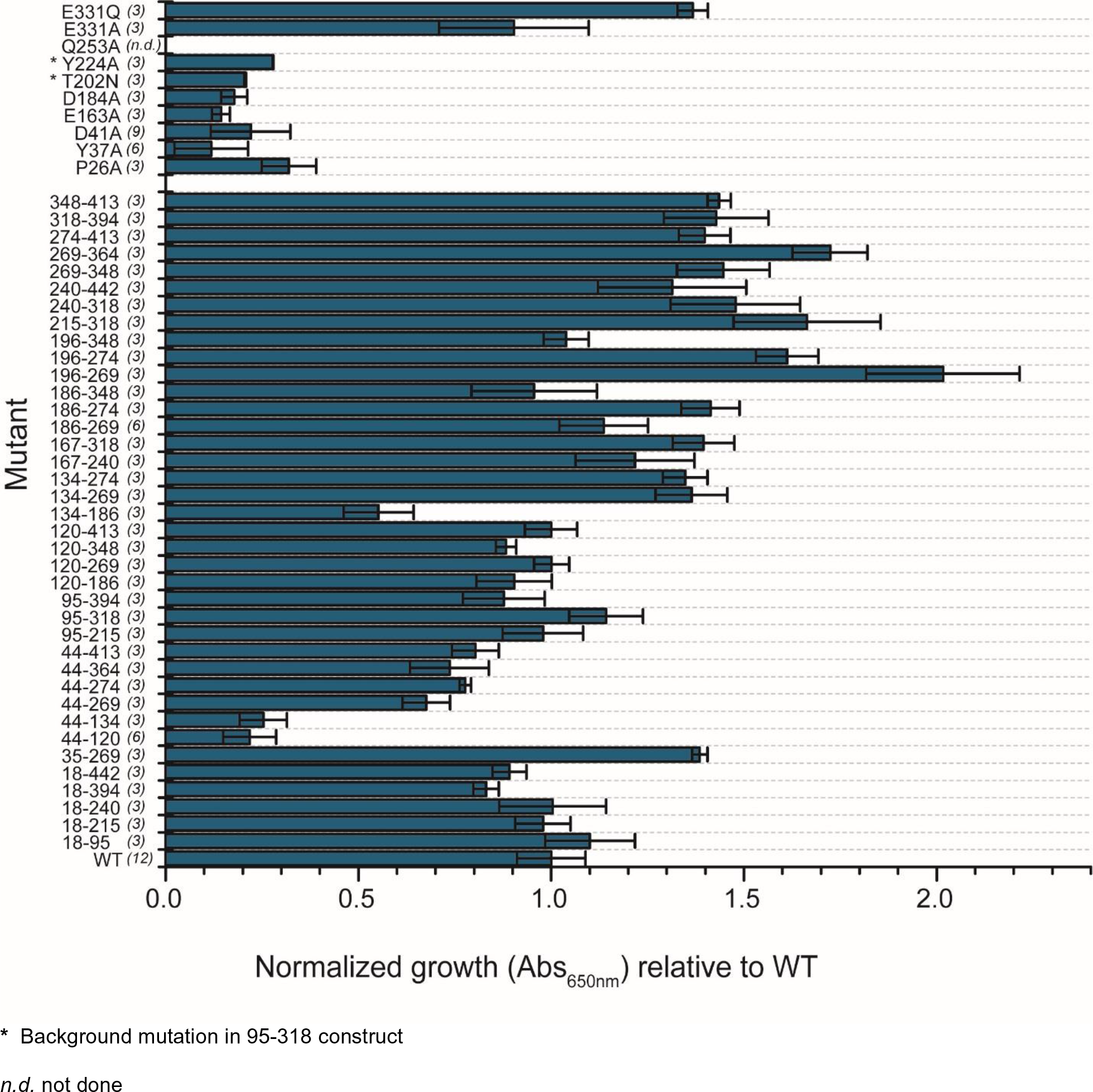
R6G resistance profiles of PfMATE mutants used for DEER spectroscopy. Growth profiles of mutants are relative to WT PfMATE after subtracting the contribution of the vector control. The bar plot highlights the average and S.E.M. for at least *n* = 3 replicates. The number of replicates is listed in brackets alongside each construct. A one-way ANOVA indicated that the population means of PfMATE mutants were significantly different at the 0.05 level: F (48, 170) = 19.73., *p* < 0.00001. A Tukey multiple comparison test showed that growth of double cysteine mutants, except 44-120 and 44-134, was not significantly different from WT, indicating that introduction of cysteines into the primary sequence of PfMATE generally has little effect on its ability to confer resistance to R6G. Conversely, growth of background mutants, except E331A/Q, was significantly different from WT, but not significantly different to the impaired mutant, D41A, indicating that these mutants are also functionally impaired.

**Supplementary Figure 2:**
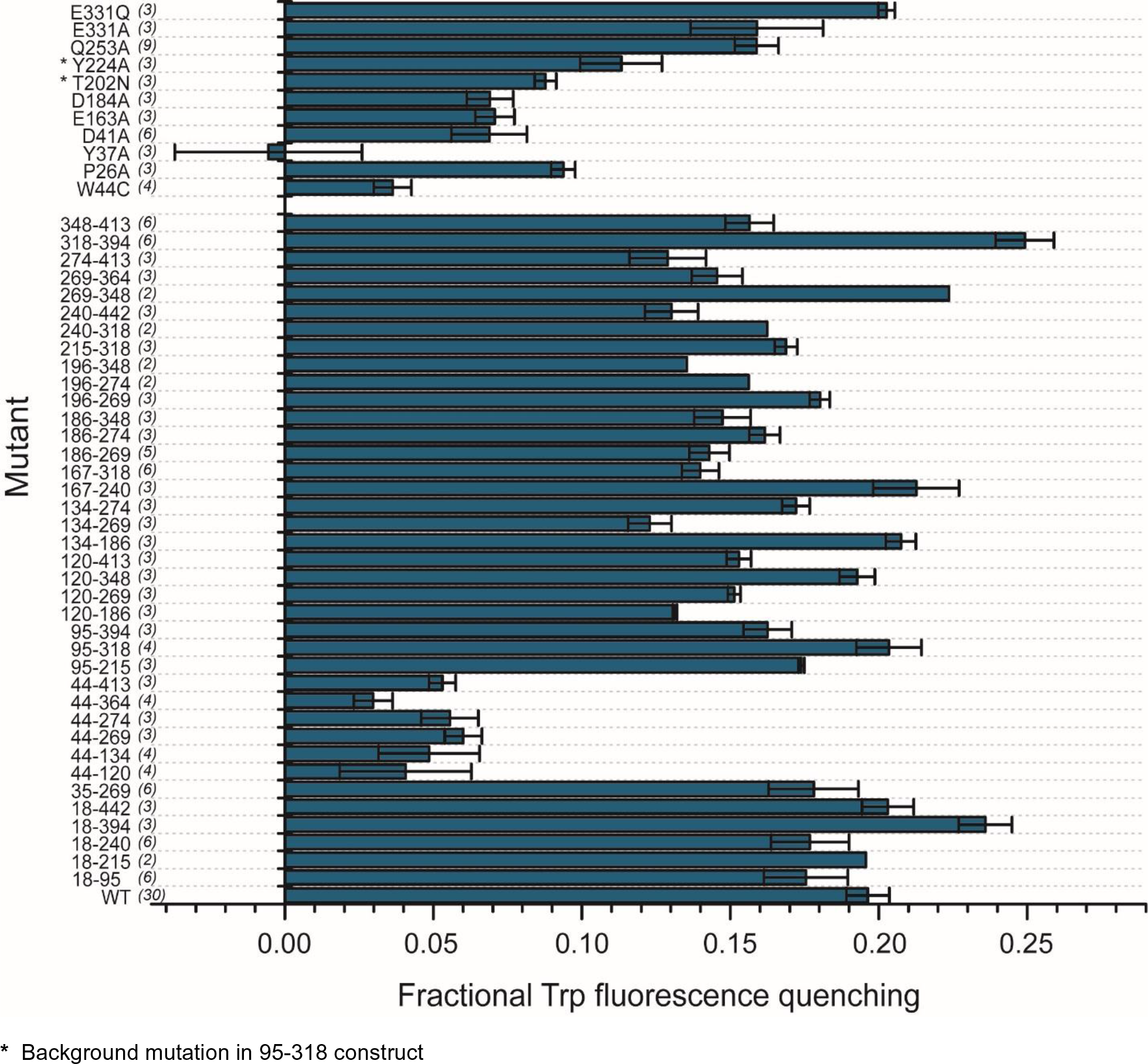
Fractional fluorescence quenching of PfMATE mutants. Fluorescence quenching at low pH due to W44 was used as a surrogate reporter of conformational changes in spin labeled PfMATE and background mutants, as detailed in methods. The number of replicates is listed in brackets alongside each construct. The bar plot highlights the average and S.E.M. for at least *n* = 3 replicates. Where *n = 2*, the average of 2 replicates are presented. For *n* > 3, data are collected from multiple protein preparations. A one-way ANOVA indicated that the population means of PfMATE mutants were significantly different at the 0.05 level: F (50, 161) = 18.80, *p* < 0.00001. A Tukey multiple comparison test showed, as expected, that double cysteine constructs containing W44C were significantly different from WT and do not demonstrate significant fluorescence quenching at low pH. Double cysteine constructs were significantly different from W44C, indicating that introduction of cysteines into the primary sequence has minimal effects on the pH sensor of PfMATE. For the background mutants (except E331A/Q) Trp quenching was not significantly different from W44C.

**Supplementary Figure 3:**
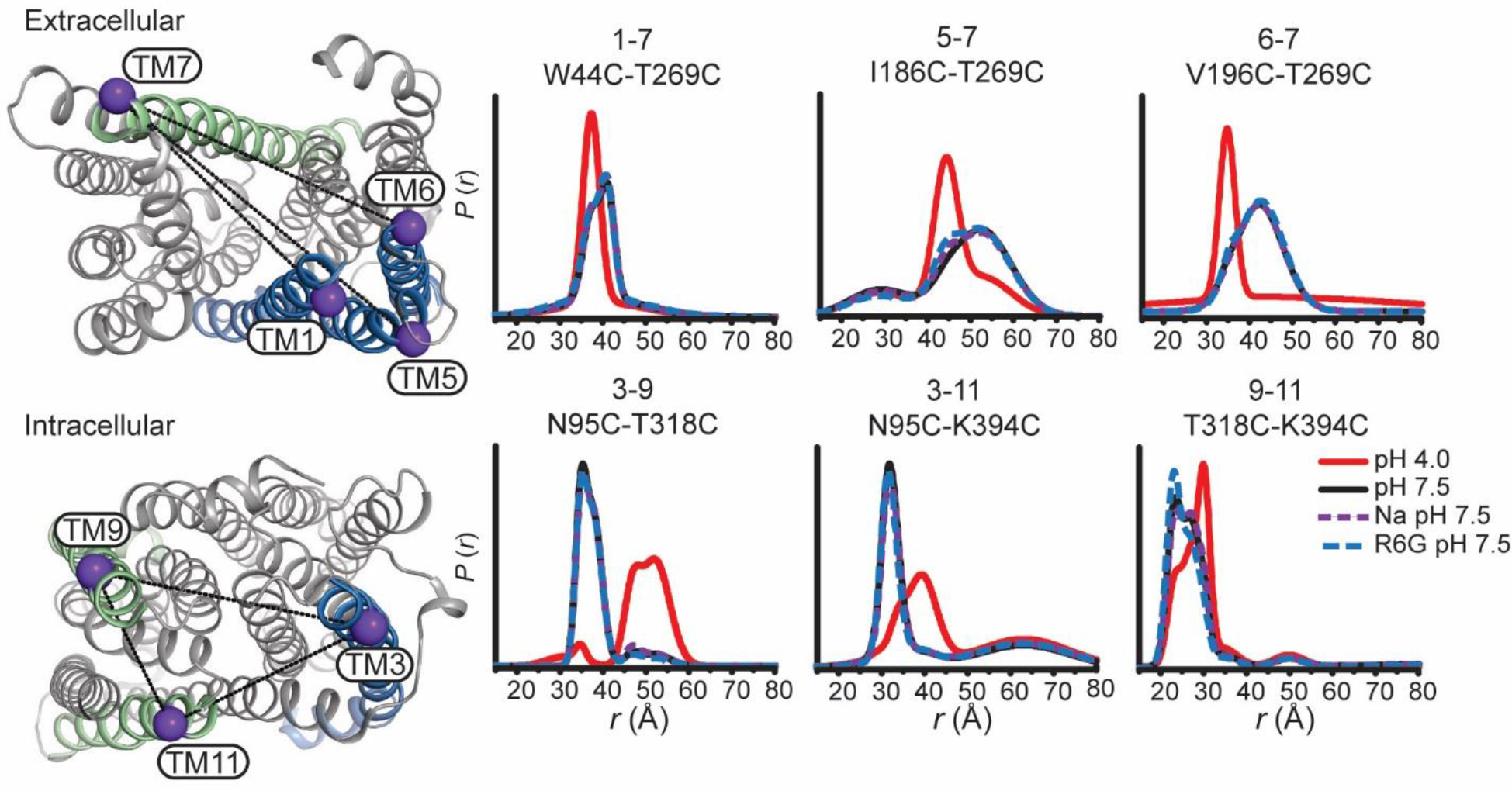
Conformational change is driven by H^+^. Sodium (dashed purple traces) and R6G (dashed blue traces) were added to a final concentration of 50 mM and 1 mM, respectively, at pH 7.5. Both ligands demonstrate limited effects on conformational change. The locations of representative spin label pairs on the extracellular (top) and intracellular sides (bottom) are highlighted on the OF structure by purple spheres connected by a line.

**Supplementary Figure 4:**
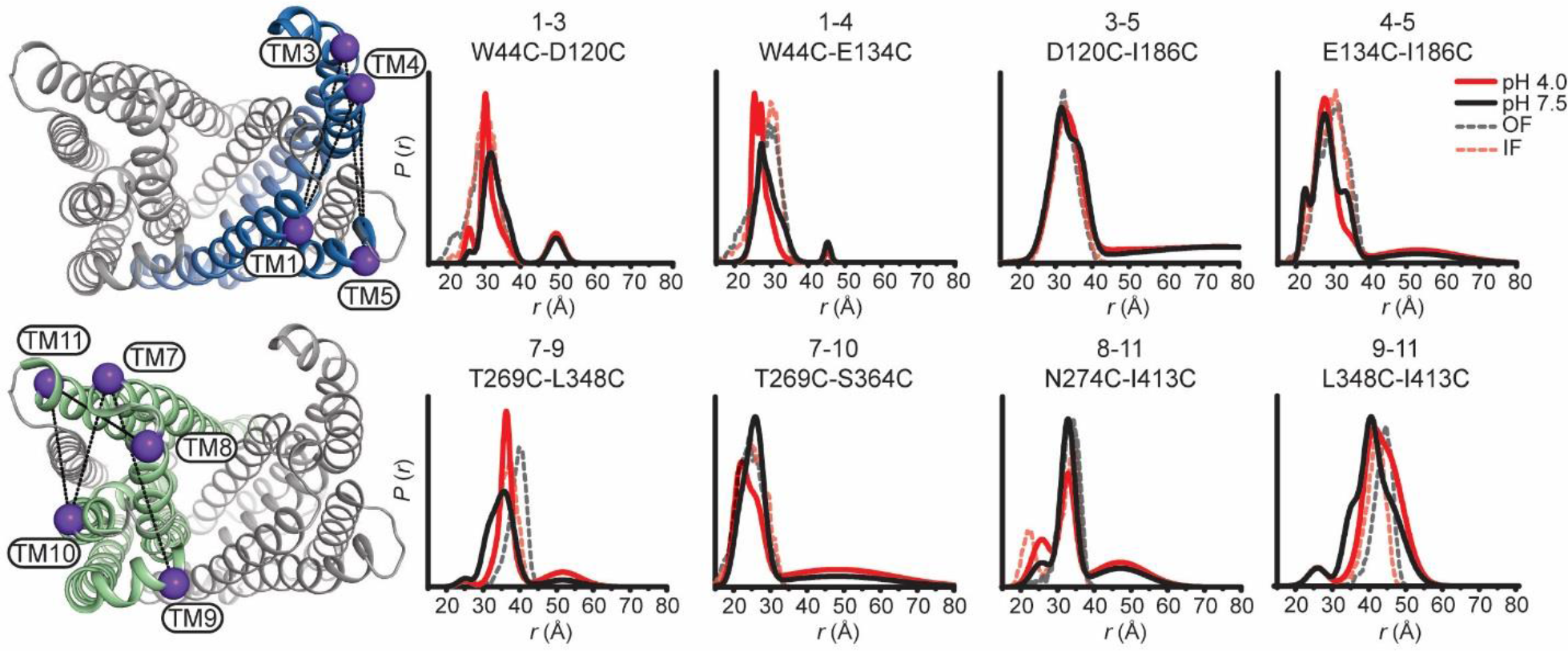
PfMATE intradomain DEER measurements on the extracellular side demonstrate limited conformational changes. Spin labeled positions on TMs in the N-lobe (top) and C-lobe (bottom) are highlighted on the OF structure by purple spheres. Experimentally-determined distributions (solid lines) are plotted with the predicted distance distributions derived from the OF (black, dashed traces) and the IF (red, dashed traces) crystal structures.

**Supplementary Figure 5:**
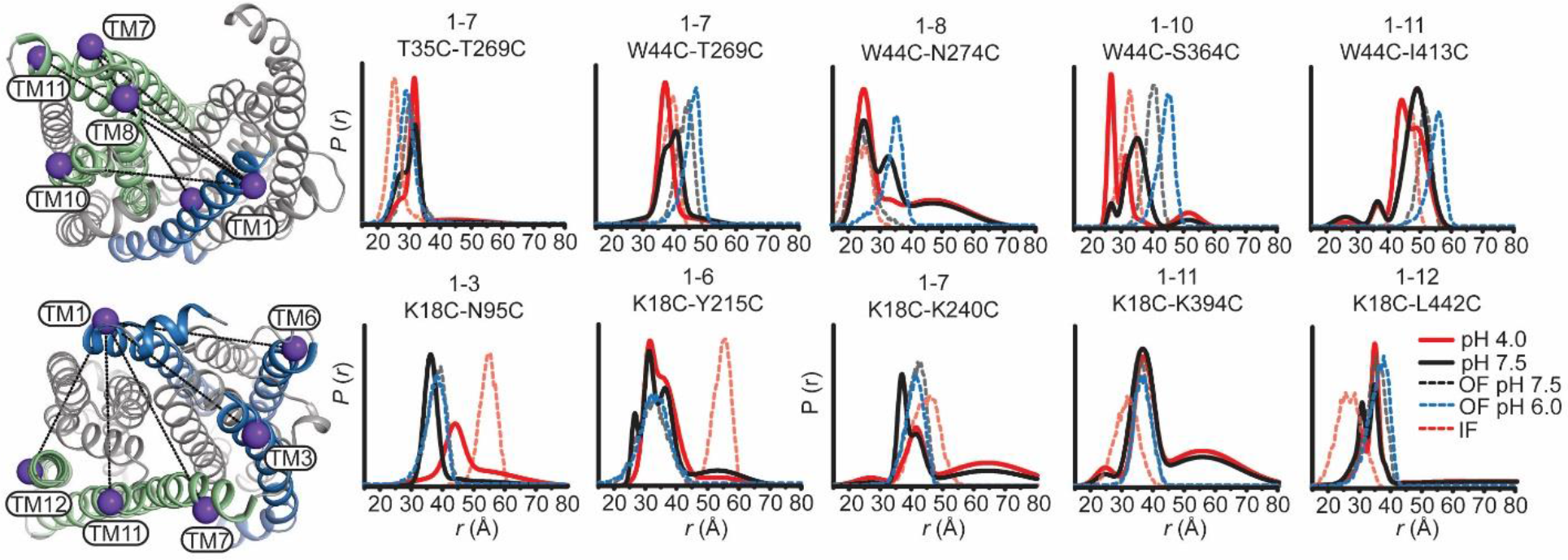
TM1 conformational changes diverge from structural models. Extracellular (top panels) and intracellular (bottom panels) measurements from TM1 are highlighted on the OF structure of PfMATE. Experimentally determined distributions from TM1 are plotted with the predicted distributions derived from the OF crystal structures obtained at pH 8.0 and 6.0 (black and blue dashed traces, respectively) and the IF (red, dashed traces) crystal structure. Conformational changes are observed mainly as population shifts to shorter distances on the extracellular side, which is inconsistent with the longer predicted distances from the OF pH 6.0 structure. On the intracellular side, limited conformational change is observed except between TM1 and TM3, consistent with the conformational changes described in the latter.

**Supplementary Figure 6:**
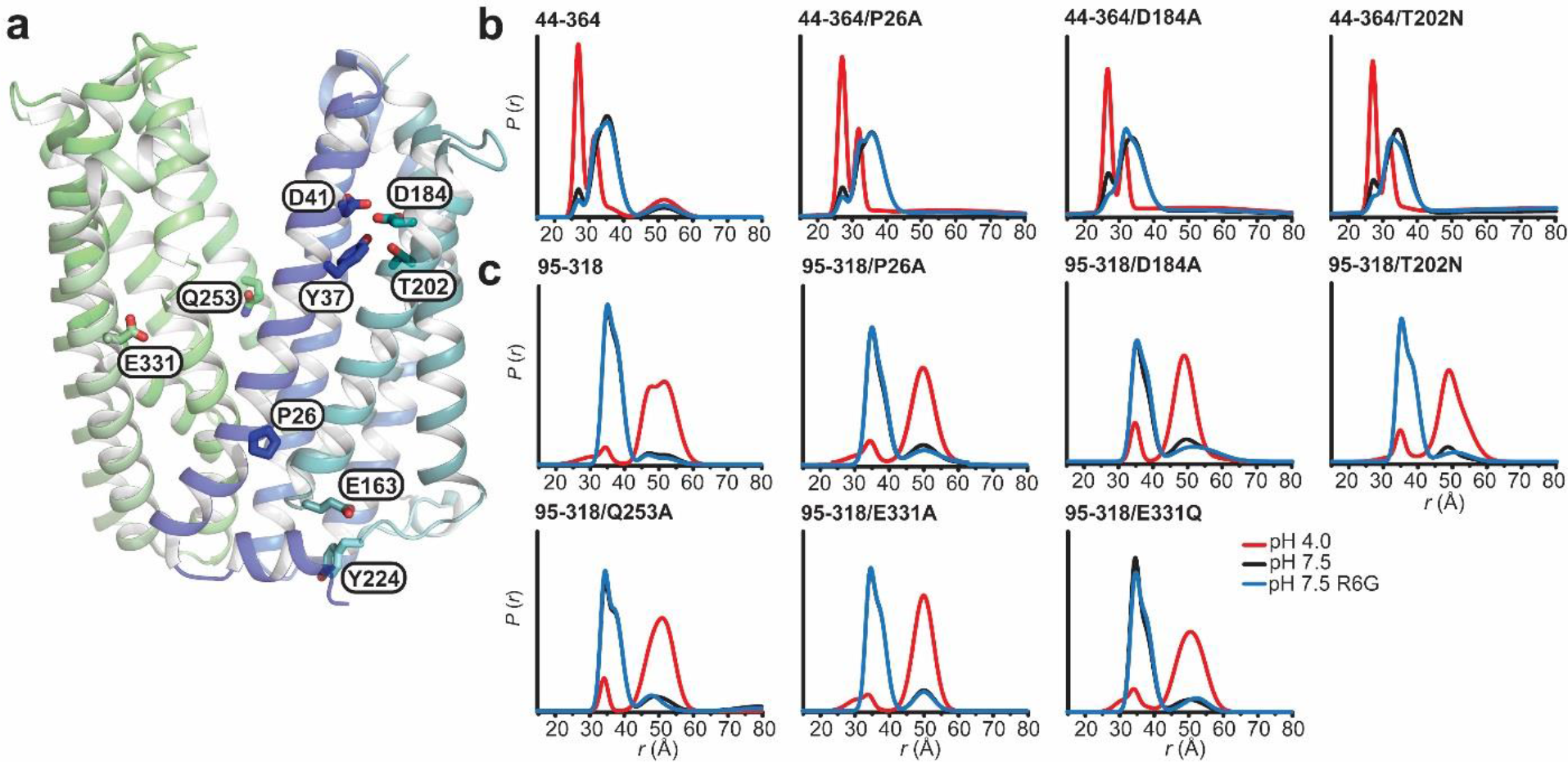
Conformational dynamics of PfMATE N-lobe mutants. (A): Sidechains of functionally essential residues in the N-lobe, as well as Q253 and E331, thought to be involved in lipid and proton binding, respectively, are depicted as sticks on the OF PfMATE structure. Mutations of these residues in reporter pairs on the extracellular (B) and intracellular (C) sides reveal that these residues do not affect the distance changes at low pH.

**Supplementary Figure 7:**
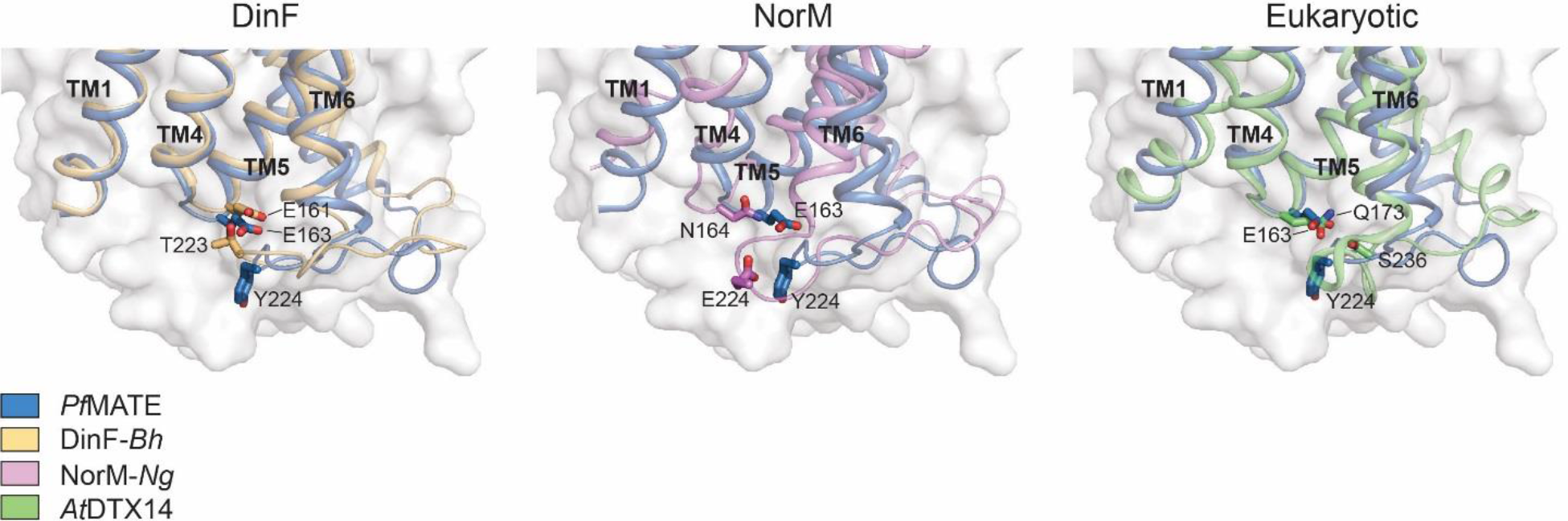
IL4-5 may be implicated in conformational switching across the MATE family of transporters. Structural alignments of PfMATE with representative MATEs from each subfamily reveal residues on the IL4-5 loop that may have H-bond or electrostatic interactions with residues on the IL6-7 loop of the transporters.

**Supplementary Table 1.**
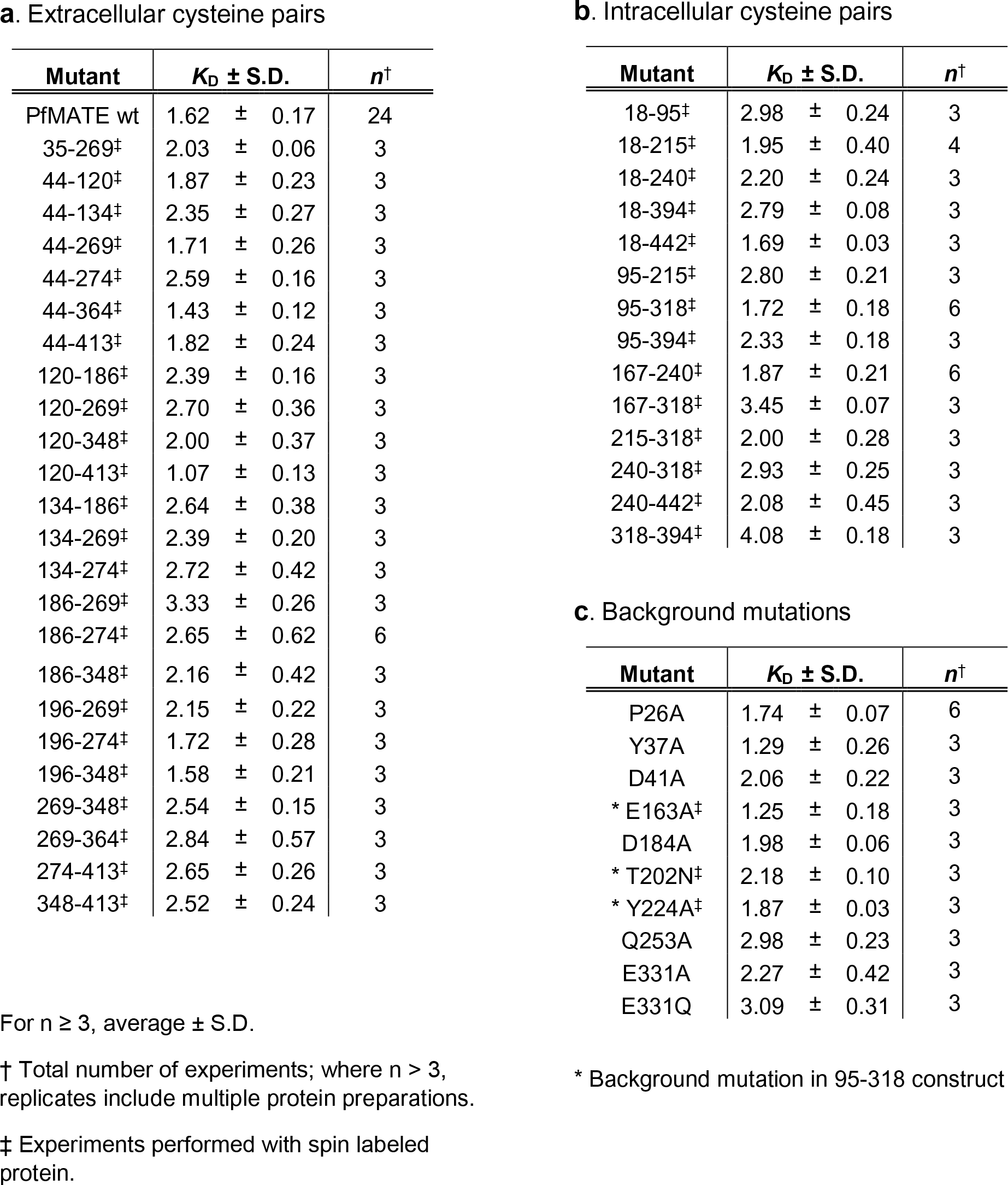

## EPR Data Appendices

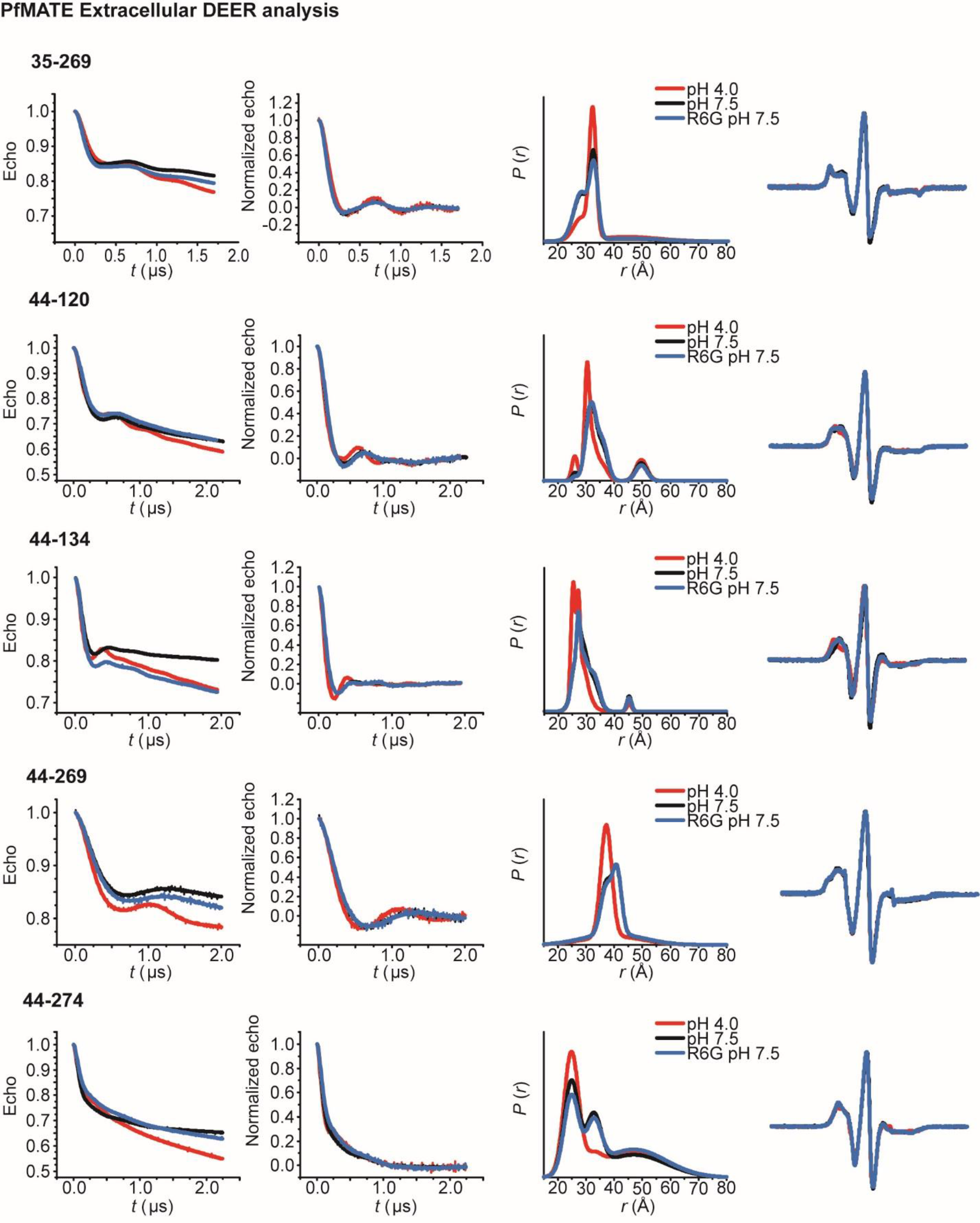
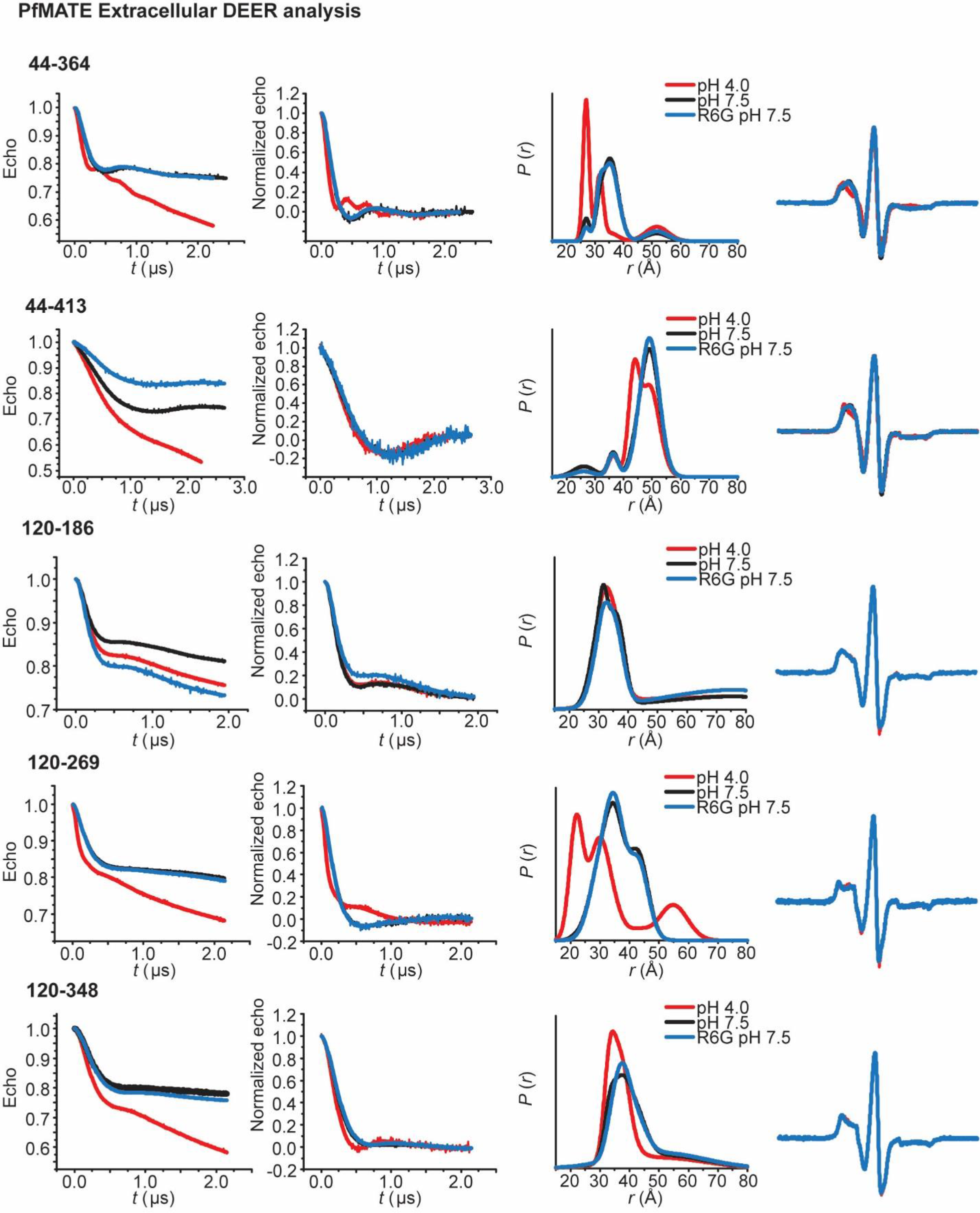
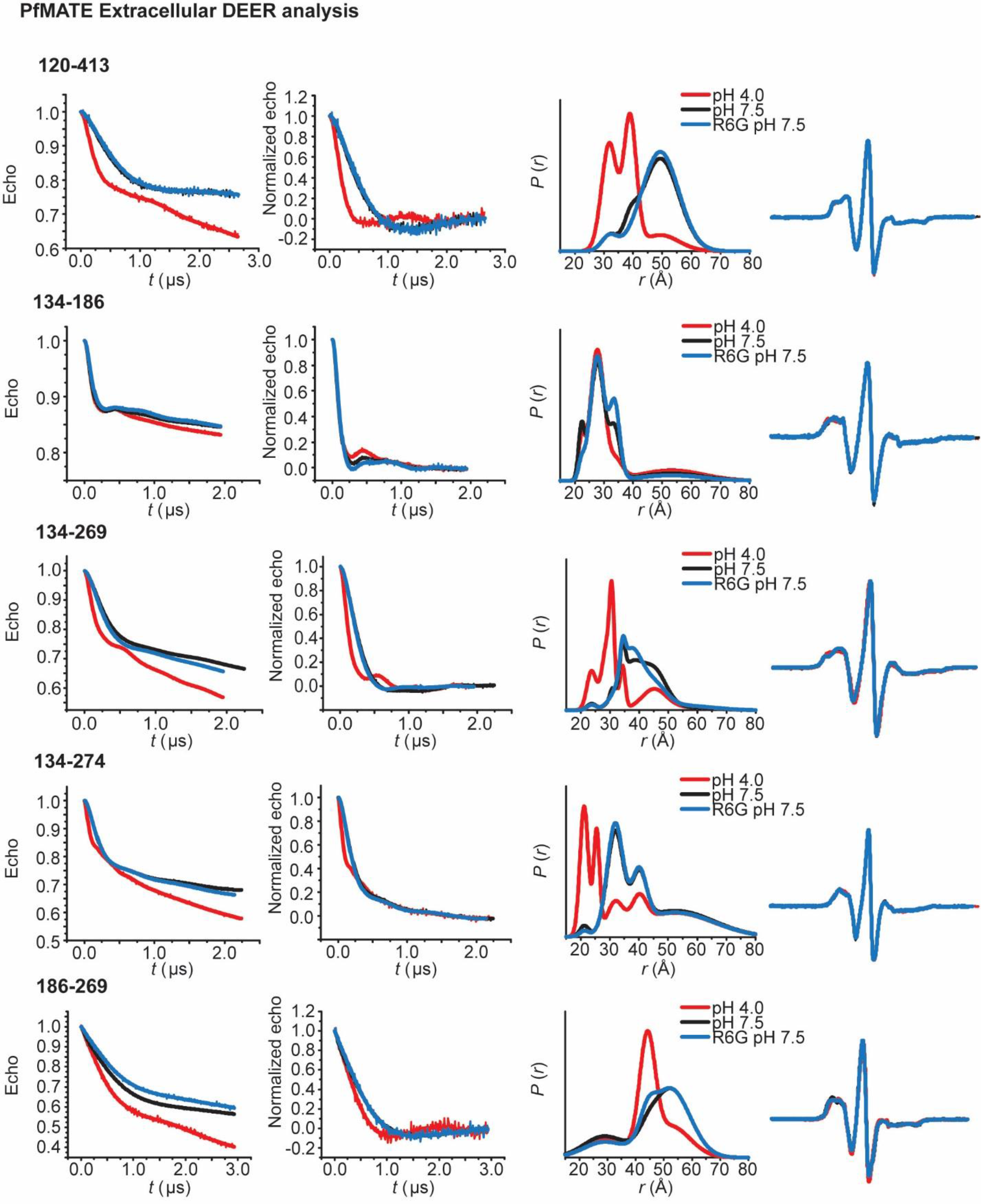
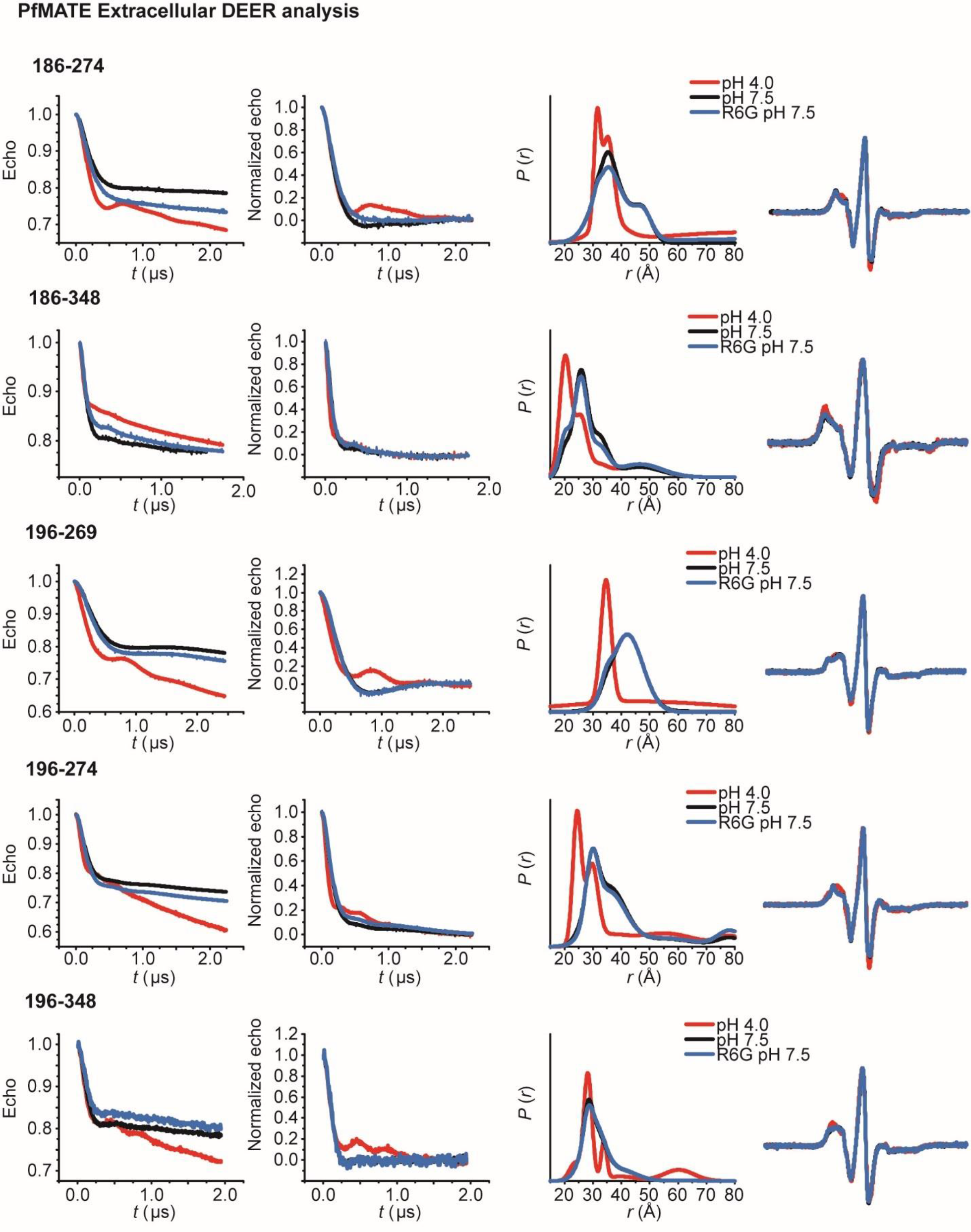
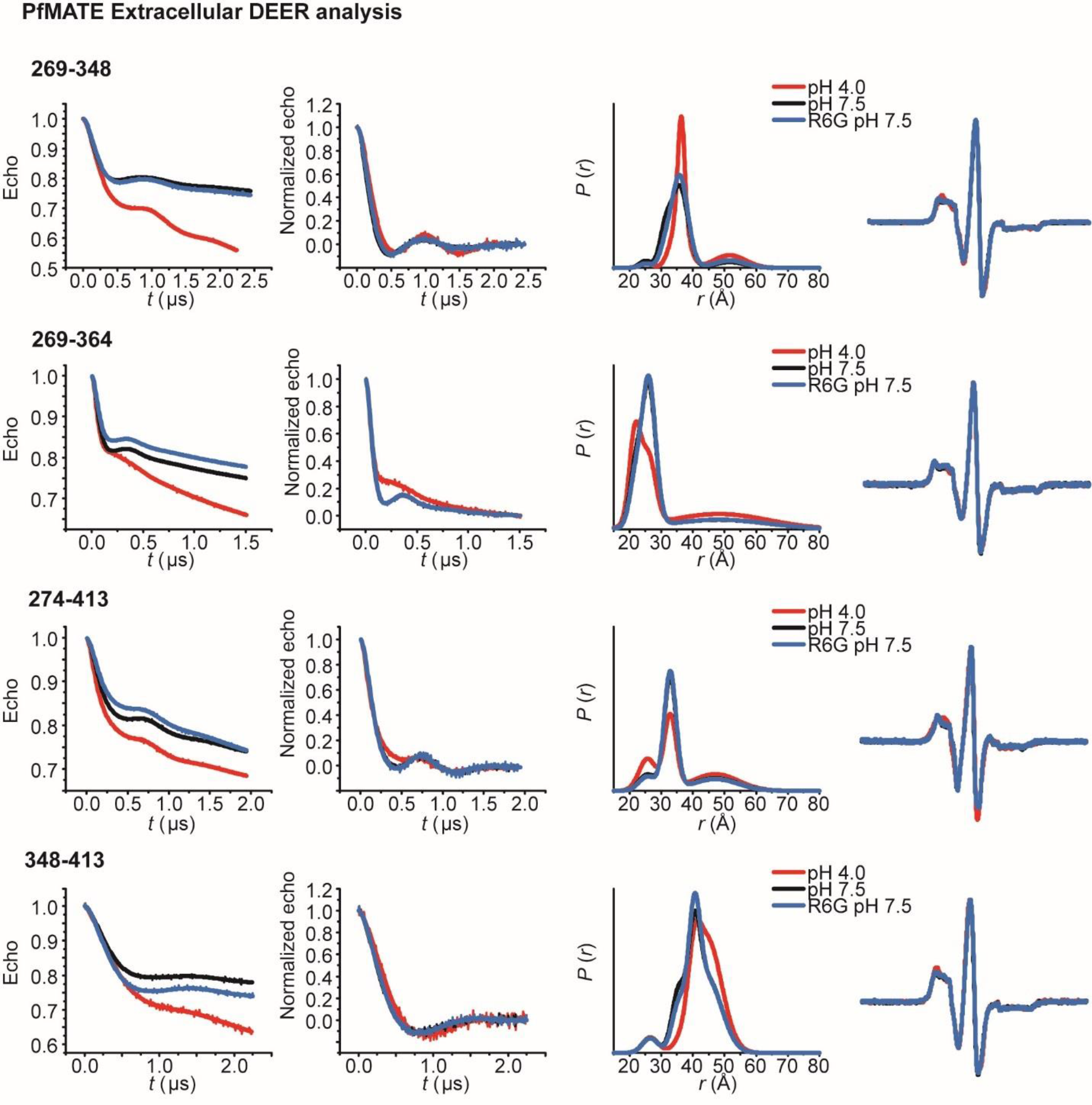
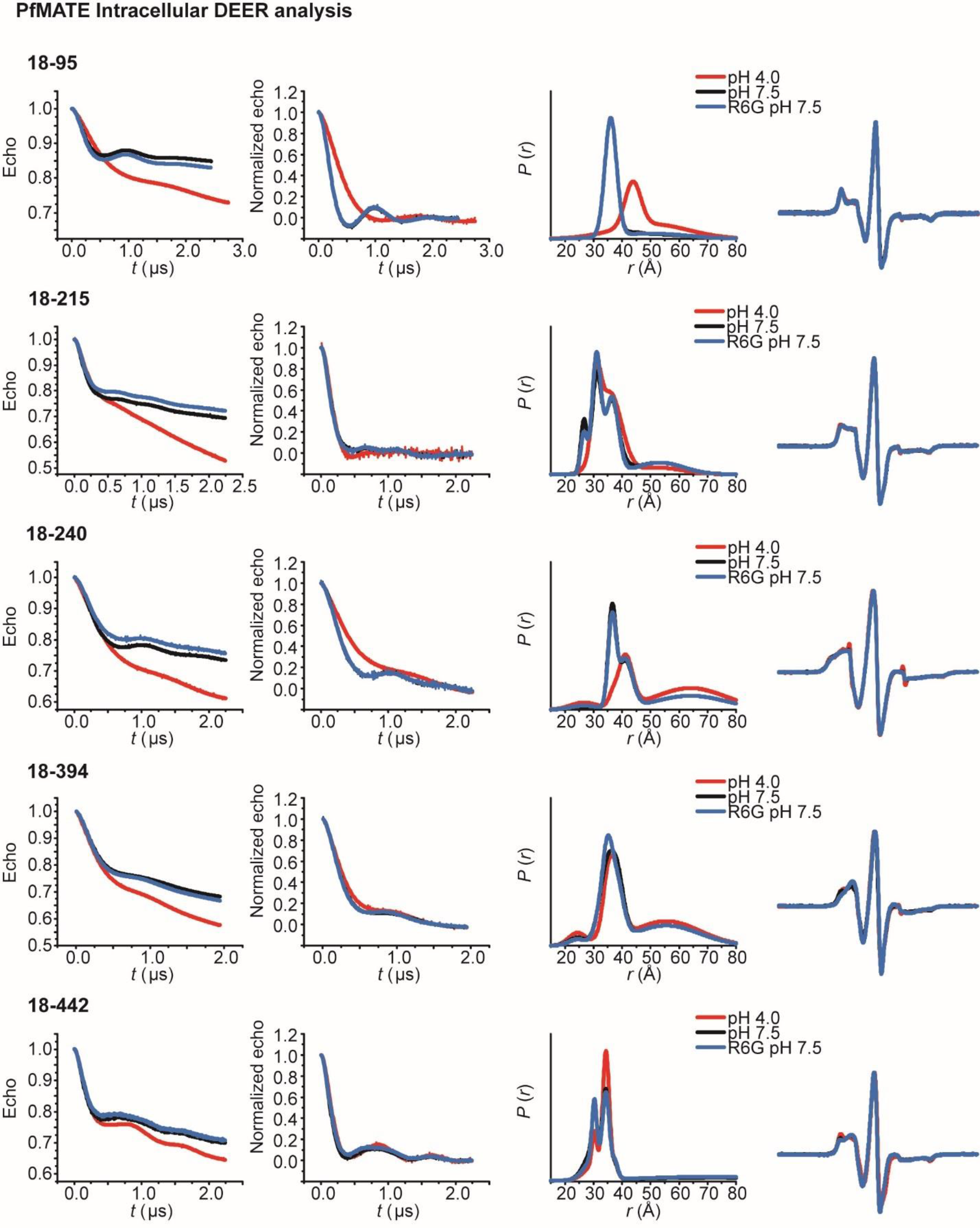
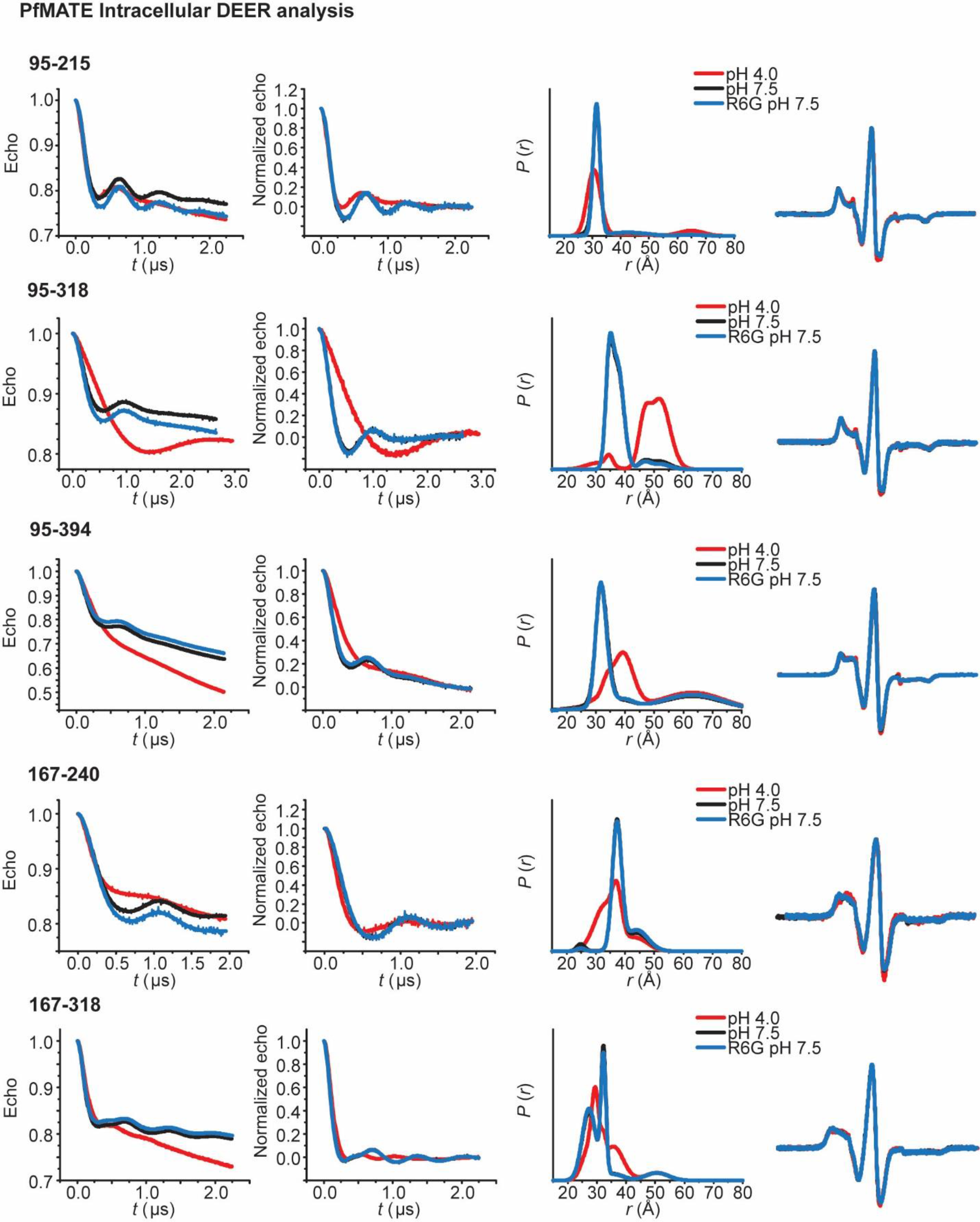
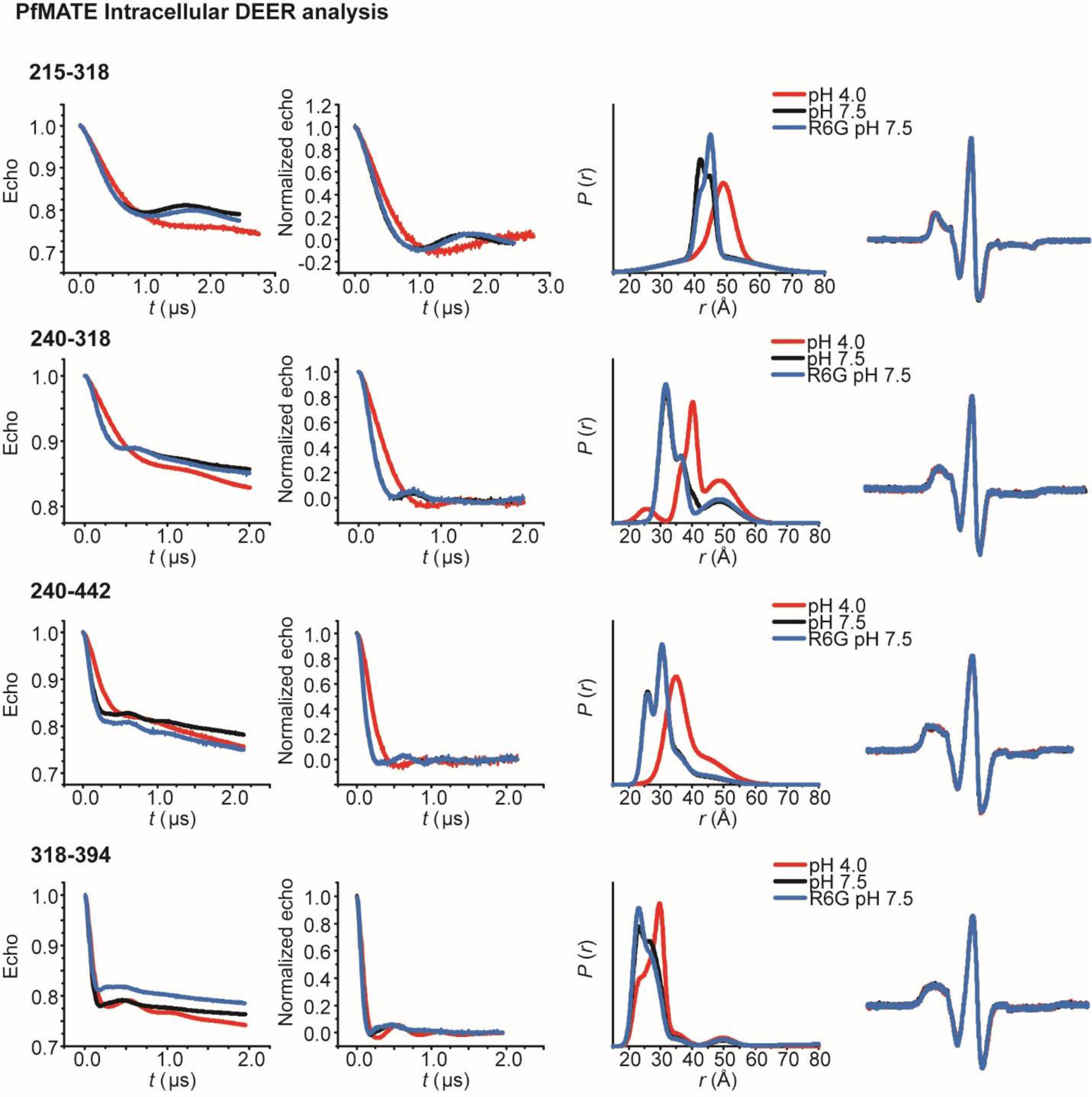
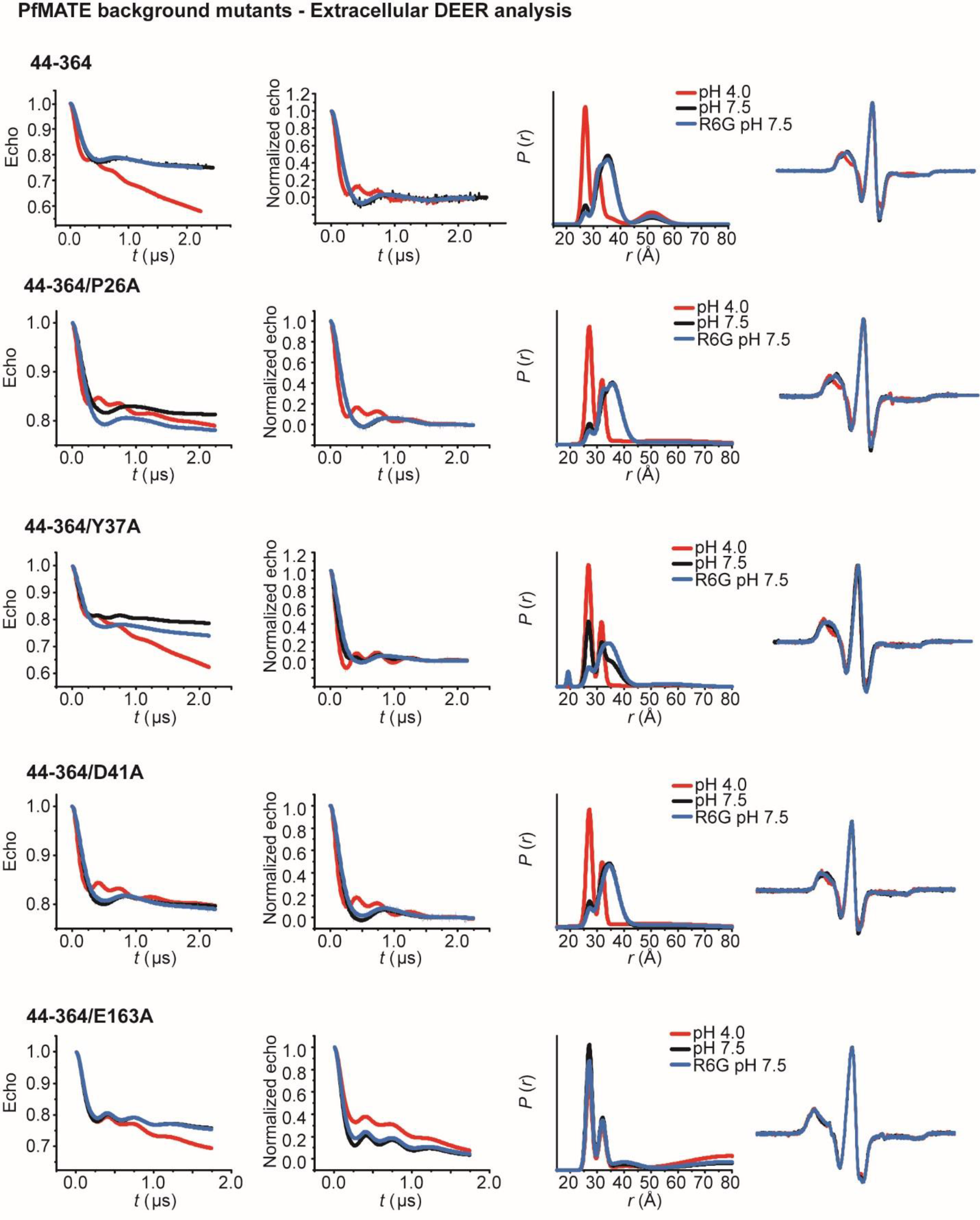
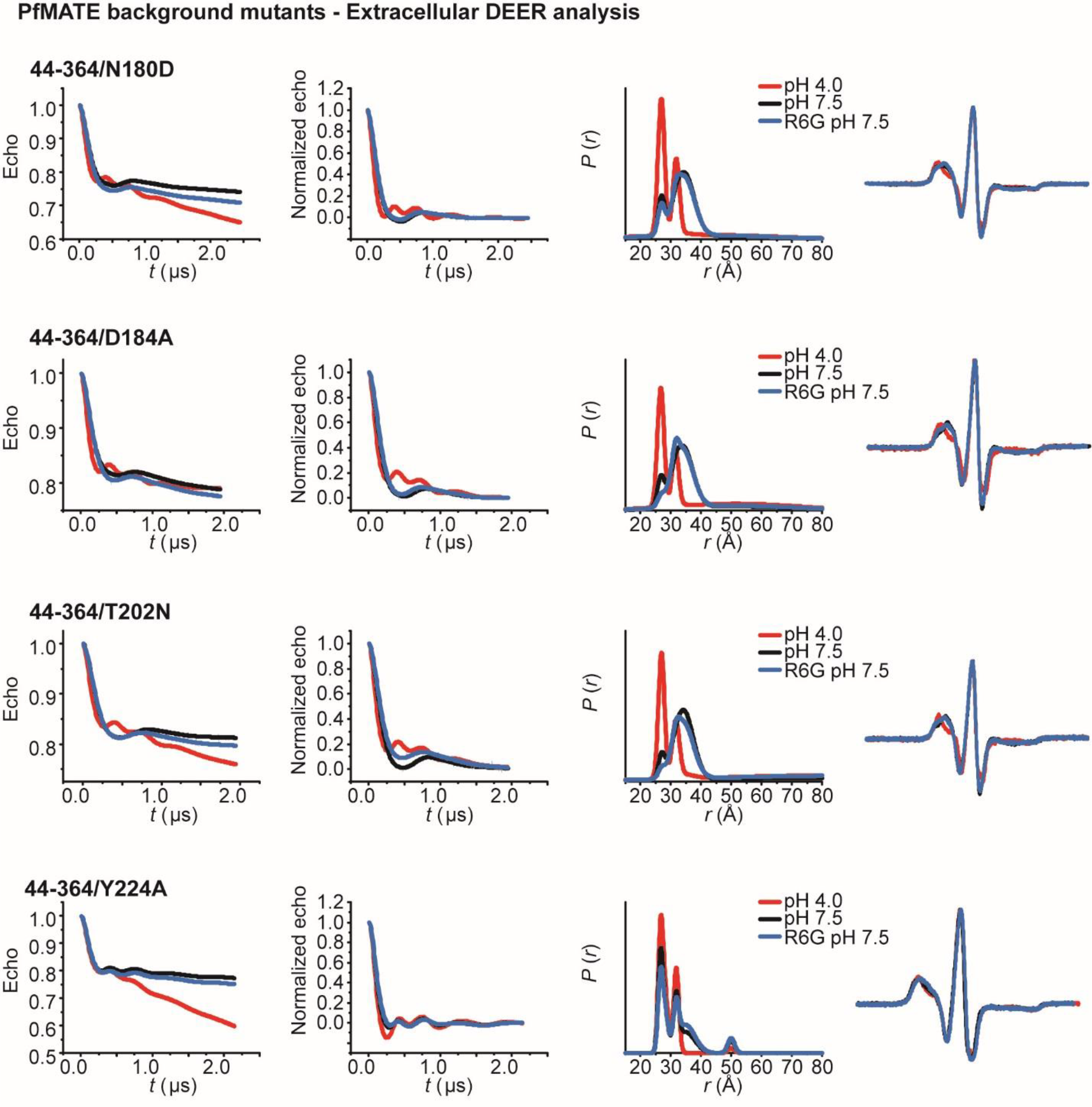
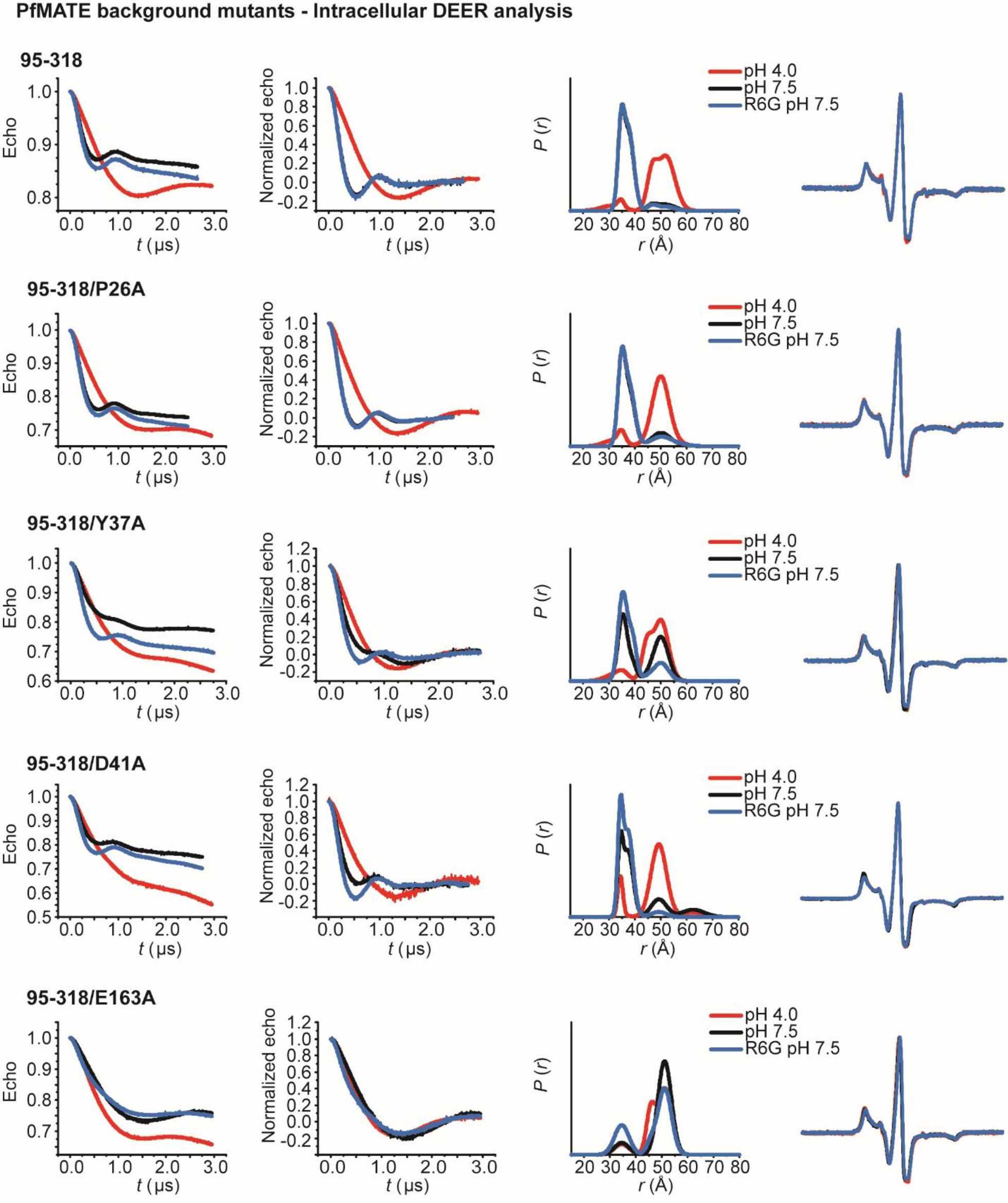
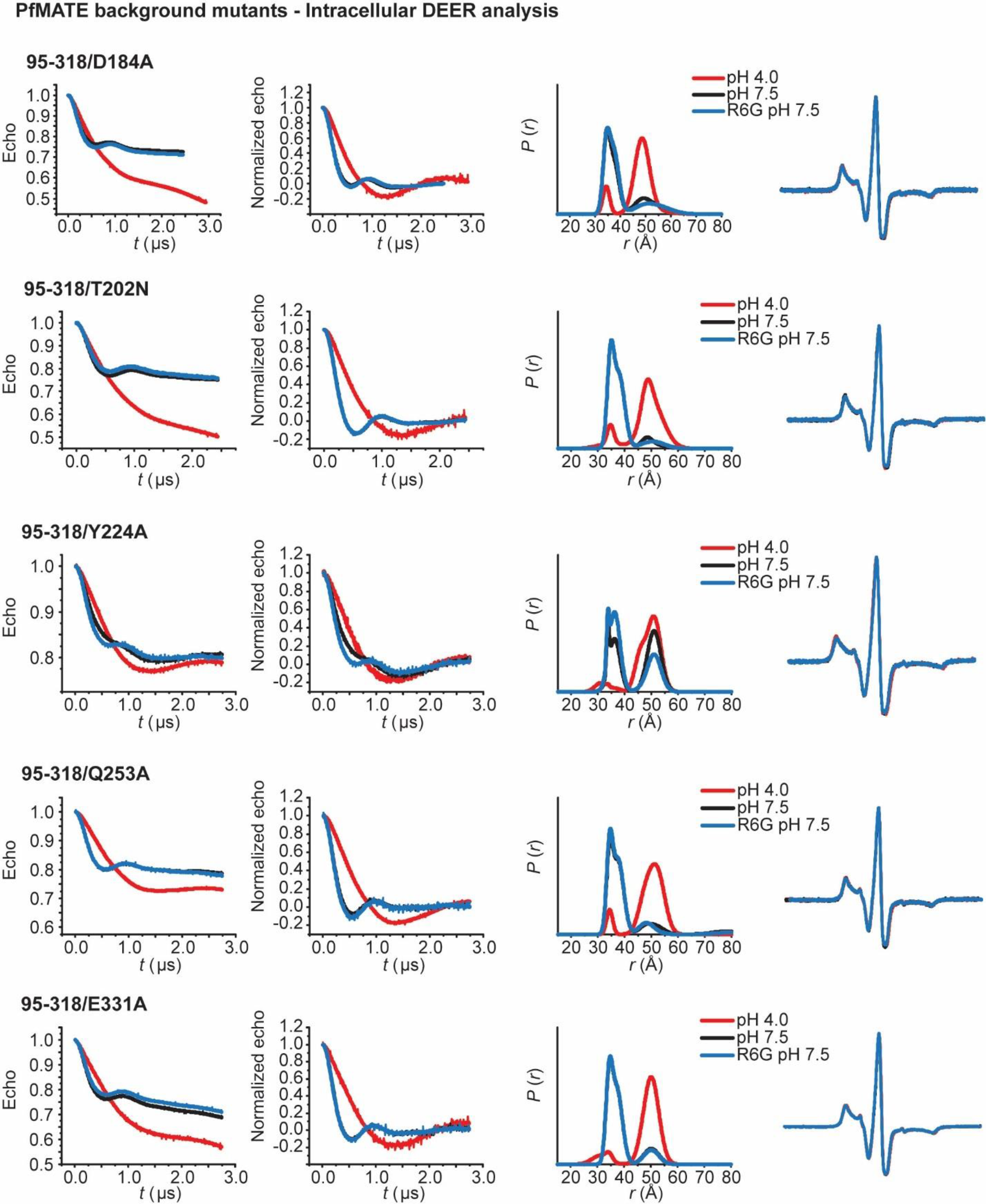
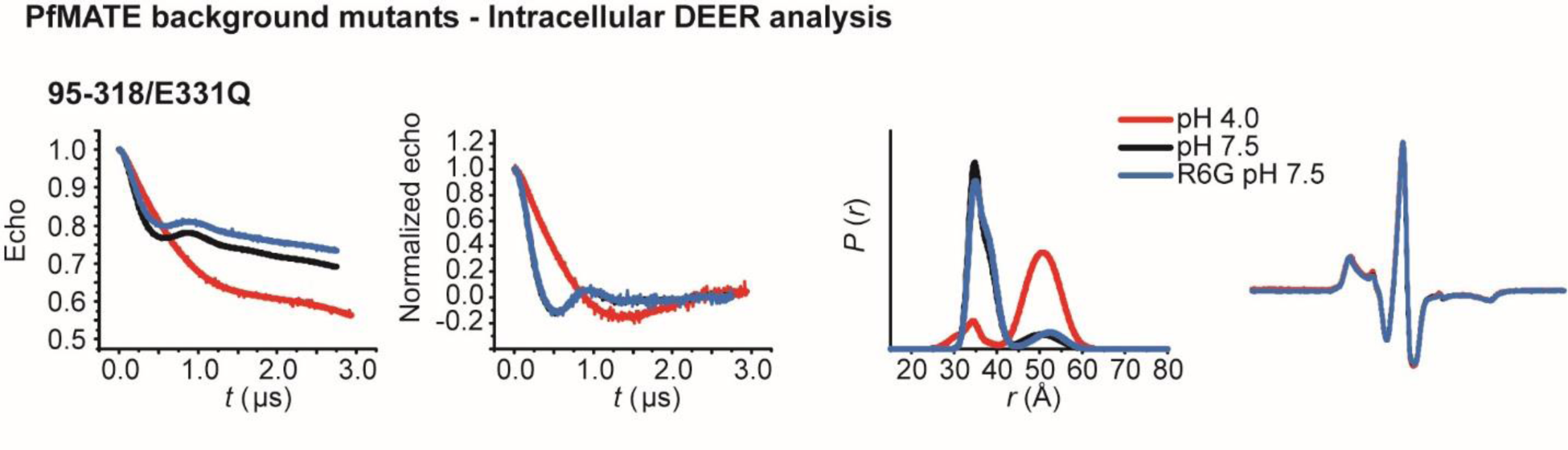

## References

1. Higgins, C. F. Multiple molecular mechanisms for multidrug resistance transporters. Nature 446, 749–757 (2007).

2. Du, D., van Veen, H. W., Murakami, S., Pos, K. M. & Luisi, B. F. Structure, mechanism and cooperation of bacterial multidrug transporters. Curr Opin Struct Biol 33, 76–91 (2015).

3. Blair, J. M., Webber, M. A., Baylay, A. J., Ogbolu, D. O. & Piddock, L. J. Molecular mechanisms of antibiotic resistance. Nat Rev Microbiol 13, 42–51 (2014).

4. Tal, N. & Schuldiner, S. A coordinated network of transporters with overlapping specificities provides a robust survival strategy. Proc Natl Acad Sci U S A 106, 9051–9056 (2009).

5. Jardetzky, O. Simple allosteric model for membrane pumps. Nature 211, 969–970 (1966).

6. Mitchell, P. A general theory of membrane transport from studies of bacteria. Nature (1957). doi:10.1038/180134a0

7. Morita, Y., Kataoka, A., Shiota, S., Mizushima, T. & Tsuchiya, T. NorM of Vibrio parahaemolyticus is an Na+-driven multidrug efflux pump. J. Bacteriol. (2000). doi:10.1128/JB.182.23.6694-6697.2000

8. Chen, J. et al. VmrA, a member of a novel class of Na(+)-coupled multidrug efflux pumps from Vibrio parahaemolyticus. J Bacteriol 184, 572–576 (2002).

9. Long, F., Rouquette-Loughlin, C., Shafer, W. M. & Yu, E. W. Functional cloning and characterization of the multidrug efflux pumps NorM from Neisseria gonorrhoeae and YdhE from Escherichia coli. Antimicrob Agents Chemother 52, 3052–3060 (2008).

10. He, G. X. et al. An H+-Coupled Multidrug Efflux Pump, PmpM, a Member of the MATE Family of Transporters, from Pseudomonas aeruginosa. J. Bacteriol. (2004). doi:10.1128/JB.186.1.262-265.2004

11. Su, X. Z., Chen, J., Mizushima, T., Kuroda, T. & Tsuchiya, T. AbeM, an H+-coupled Acinetobacter baumannii multidrug efflux pump belonging to the MATE family of transporters. Antimicrob Agents Chemother 49, 4362–4364 (2005).

12. Li, L., He, Z., Pandey, G. K., Tsuchiya, T. & Luan, S. Functional cloning and characterization of a plant efflux carrier for multidrug and heavy metal detoxification. J. Biol. Chem. (2002). doi:10.1074/jbc.M108777200

13. Masuda, S. et al. Identification and functional characterization of a new human kidney-specific H+/organic cation antiporter, kidney-specific multidrug and toxin extrusion 2. J Am Soc Nephrol 17, 2127–2135 (2006).

14. Lu, M. Structures of multidrug and toxic compound extrusion transporters and their mechanistic implications. Channels (Austin) 10, 88–100 (2016).

15. Hvorup, R. N. et al. The multidrug/oligosaccharidyl-lipid/polysaccharide (MOP) exporter superfamily. European Journal of Biochemistry (2003). doi:10.1046/j.1432-1033.2003.03418.x

16. He, X. et al. Structure of a cation-bound multidrug and toxic compound extrusion transporter. Nature 467, 991–994 (2010).

17. Lu, M., Radchenko, M., Symersky, J., Nie, R. & Guo, Y. Structural insights into H+-coupled multidrug extrusion by a MATE transporter. Nat Struct Mol Biol 20, 1310–1317 (2013).

18. Lu, M. et al. Structures of a Na+-coupled, substrate-bound MATE multidrug transporter. Proc Natl Acad Sci U S A 110, 2099–2104 (2013).

19. Tanaka, Y. et al. Structural basis for the drug extrusion mechanism by a MATE multidrug transporter. Nature 496, 247–51 (2013).

20. Mousa, J. J., Newsome, R. C., Yang, Y., Jobin, C. & Bruner, S. D. ClbM is a versatile, cation-promiscuous MATE transporter found in the colibactin biosynthetic gene cluster. Biochem Biophys Res Commun 482, 1233–1239 (2016).

21. Miyauchi, H. et al. Structural basis for xenobiotic extrusion by eukaryotic MATE transporter. Nat Commun 8, 1633 (2017).

22. Tanaka, Y., Iwaki, S. & Tsukazaki, T. Crystal Structure of a Plant Multidrug and Toxic Compound Extrusion Family Protein. Structure 25, 1455–1460 e2 (2017).

23. Kusakizako, T. et al. Structural Basis of H(+)-Dependent Conformational Change in a Bacterial MATE Transporter. Structure 27, 293–301 e3 (2019).

24. Steed, P. R., Stein, R. A., Mishra, S., Goodman, M. C. & Mchaourab, H. S. Na(+)-substrate coupling in the multidrug antiporter norm probed with a spin-labeled substrate. Biochemistry 52, 5790–5799 (2013).

25. Zakrzewska, S. et al. Inward-facing conformation of a multidrug resistance MATE family transporter. Proc Natl Acad Sci U S A 116, 12275–12284 (2019).

26. Jagessar, K. L., Mchaourab, H. S. & Claxton, D. P. The N-terminal domain of an archaeal multidrug and toxin extrusion (MATE) transporter mediates proton coupling required for prokaryotic drug resistance. J Biol Chem (2019).

27. Ficici, E., Zhou, W., Castellano, S. & Faraldo-Gómez, J. D. Broadly conserved Na + – binding site in the N-lobe of prokaryotic multidrug MATE transporters. Proc. Natl. Acad. Sci. 115, E6172–E6181 (2018).

28. Dastvan, R., Mishra, S., Peskova, Y. B., Nakamoto, R. K. & Mchaourab, H. S. Mechanism of allosteric modulation of P-glycoprotein by transport substrates and inhibitors. Science (80-.). (2019). doi:10.1126/science.aav9406

29. Claxton, D. P., Kazmier, K., Mishra, S. & Mchaourab, H. S. Navigating Membrane Protein Structure, Dynamics, and Energy Landscapes Using Spin Labeling and EPR Spectroscopy. Methods Enzym. 564, 349–387 (2015).

30. Jeschke, G. DEER distance measurements on proteins. Annu Rev Phys Chem 63, 419–446 (2012).

31. Mchaourab, H. S., Steed, P. R. & Kazmier, K. Toward the fourth dimension of membrane protein structure: insight into dynamics from spin-labeling EPR spectroscopy. Structure 19, 1549–1561 (2011).

32. Schiemann, O. & Prisner, T. F. Long-range distance determinations in biomacromolecules by EPR spectroscopy. Q Rev Biophys 40, 1–53 (2007).

33. Claxton, D. P., Jagessar, K. L., Steed, P. R., Stein, R. A. & Mchaourab, H. S. Sodium and proton coupling in the conformational cycle of a MATE antiporter from Vibrio cholerae. Proc. Natl. Acad. Sci. (2018). doi:10.1073/pnas.1802417115

34. Tanaka, Y. et al. Structural basis for the drug extrusion mechanism by a MATE multidrug transporter. Nature 496, 247–251 (2013).

35. Lu, M., Radchenko, M., Symersky, J., Nie, R. & Guo, Y. Structural insights into H+-coupled multidrug extrusion by a MATE transporter. Nat. Struct. Mol. Biol. 20, 1310–7 (2013).

36. Ficici, E., Zhou, W., Castellano, S. & Faraldo-Gomez, J. D. Broadly conserved Na(+)-binding site in the N-lobe of prokaryotic multidrug MATE transporters. Proc Natl Acad Sci U S A 115, E6172–E6181 (2018).

37. Jo, S., Kim, T., Iyer, V. G. & Im, W. CHARMM-GUI: a web-based graphical user interface for CHARMM. J Comput Chem 29, 1859–1865 (2008).

38. Kuk, A. C. Y., Hao, A., Guan, Z. & Lee, S. Y. Visualizing conformation transitions of the Lipid II flippase MurJ. Nat. Commun. 10, (2019).

39. Eisinger, M. L., Nie, L., Dörrbaum, A. R., Langer, J. D. & Michel, H. The Xenobiotic Extrusion Mechanism of the MATE Transporter NorM_PS from Pseudomonas stutzeri. J. Mol. Biol. 430, 1311–1323 (2018).

40. Singh, A. K., Haldar, R., Mandal, D. & Kundu, M. Analysis of the topology of Vibrio cholerae NorM and identification of amino acid residues involved in norfloxacin resistance. Antimicrob Agents Chemother 50, 3717–3723 (2006).

41. Nie, L. et al. Identification of the High-affinity Substrate-binding Site of the Multidrug and Toxic Compound Extrusion (MATE) Family Transporter from Pseudomonas stutzeri. J Biol Chem 291, 15503–15514 (2016).

42. Otsuka, M. et al. Identification of essential amino acid residues of the NorM Na+/multidrug antiporter in Vibrio parahaemolyticus. J Bacteriol 187, 1552–1558 (2005).

43. Jin, Y., Nair, A. & Van Veen, H. W. Multidrug transport protein NorM from Vibrio cholerae simultaneously couples to sodium- and proton-motive force. J. Biol. Chem. (2014). doi:10.1074/jbc.M113.546770

44. Song, J., Ji, C. & Zhang, J. Z. Insights on Na(+) binding and conformational dynamics in multidrug and toxic compound extrusion transporter NorM. Proteins 82, 240–249 (2013).

45. Masureel, M. et al. Protonation drives the conformational switch in the multidrug transporter LmrP. Nat. Chem. Biol. (2014). doi:10.1038/nchembio.1408

46. Vanni, S., Campomanes, P., Marcia, M. & Rothlisberger, U. Ion binding and internal hydration in the multidrug resistance secondary active transporter NorM investigated by molecular dynamics simulations. Biochemistry 51, 1281–1287 (2012).

47. Celniker, G. et al. ConSurf: Using evolutionary data to raise testable hypotheses about protein function. Israel Journal of Chemistry (2013). doi:10.1002/ijch.201200096

48. Ashkenazy, H., et al. ConSurf 2016: an improved methodology to estimate and visualize evolutionary conservation in macromolecules. Nucleic Acids Res. (2016). doi:10.1093/nar/gkw408

49. Mishra, S. et al. Conformational dynamics of the nucleotide binding domains and the power stroke of a heterodimeric ABC transporter. Elife 3, e02740 (2014).

50. Zou, P. & Mchaourab, H. S. Increased sensitivity and extended range of distance measurements in Spin-labeled membrane proteins: Q-band double electron-electron resonance and nanoscale bilayers. Biophys. J. (2010). doi:10.1016/j.bpj.2009.12.4193

51. Stein, R. A., Beth, A. H. & Hustedt, E. J. A Straightforward Approach to the Analysis of Double Electron-Electron Resonance Data. Methods Enzym. 563, 531–567 (2015).

52. Zehentbauer, F. M. et al. Fluorescence spectroscopy of Rhodamine 6G: Concentration and solvent effects. Spectrochim. Acta – Part A Mol. Biomol. Spectrosc. (2014). doi:10.1016/j.saa.2013.10.062

